# Fast enzymatic HCO_3_^-^ dehydration supports photosynthetic water oxidation in Photosystem II from pea

**DOI:** 10.1101/2021.09.30.462629

**Authors:** Alexandr V. Shitov, Vasily V. Terentyev, Govindjee Govindjee

## Abstract

Carbonic anhydrase (CA) activity, associated with Photosystem II (PSII) from *Pisum sativum*, has been shown to enhance water oxidation. But, the nature of the CA activity, its origin and role in photochemistry has been under debate, since the rates of CA reactions, measured earlier, were less than the rates of photochemical reactions. Here, we demonstrate high CA activity in PSII from *Pisum sativum*, measured by HCO_3_^-^ dehydration at pH 6.5 (i.e. under optimal condition for PSII photochemistry), with kinetic parameters K_m_ of 2.7 mM; V_max_ of 2.74·10^-2^ mM·sec^-1^; k_cat_ of 1.16·10^3^ sec^-1^ and k_cat_/K_m_ of 4.1·10^5^ M^-1^ sec^-1^, showing the enzymatic nature of this activity, which k_cat_ exceeds by ∼13 times the rate of PSII, as measured by O_2_ evolution. The similar dependence of HCO_3_^-^ dehydration, of the maximal quantum yield of photochemical reactions and of O_2_ evolution on the ratio of chlorophyll/photochemical reaction center II demonstrate the interconnection of these processes on the electron donor side of PSII. Since the removal of protons is critical for fast water oxidation, and since HCO_3_^-^ dehydration consumes a proton, we suggest that CA activity, catalyzing very fast removal of protons, supports efficient water oxidation in PSII and, thus, photosynthesis in general.

## 1. Introduction

Photosystem II (PSII) is a unique pigment-protein complex, capable of using the energy of the sunlight for water oxidation accompanied by the release of molecular oxygen, protons and electrons (Wydrzynski et al., 2005; Shevela et al., 2021). The role of bicarbonate (HCO_3_^-^) in PSII function has been extensively studied over the past five decades (for references, see reviews (van Rensen et al., 1999; van Rensen and Klimov, 2005; Shevela et al., 2012)). Bicarbonate has been shown to affect the electron transfer process on both the acceptor and the donor sides of PSII (Stemler and Govindjee, 1973; Wydrzynski and Govindjee, 1975; Eaton-Rye and Govindjee, 1988; Klimov et al., 1995a, 1995b; Allakhverdiev et al., 1997; Klimov et al., 1997a, 1997b). The following reaction is known to be catalyzed by carbonic anhydrase (Moroney et al., 2011):

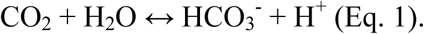

The forward reaction is CO_2_ hydration:

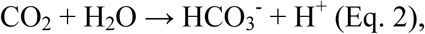

whereas, the reverse reaction is HCO_3_^-^ dehydration:

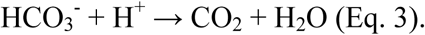

Stemler and Jursinic (Stemler and Jursinic, 1983) proposed that the rate of CO_2_ hydration by CA (Eq. 2) must be high enough for providing enough bicarbonate to PSII. This CA activity with its unique property has been found to be associated with PSII preparations, isolated from higher plants (Stemler, 1986; Pronina et al., 2002; Moskvin et al., 2004; Lu et al., 2005; Rudenko et al., 2007; Ignatova et al., 2011; Shitov et al., 2009; Hillier et al., 2006). However, this hydration activity (in PSII) was found to be rather low (12-15 Wilbur – Anderson units (W-A un.)·mg^-1^ of Chlorophyll (Chl) (Shitov et al., 2018, 2009)) and the results of dehydration activity (Eq. 3) measurements (Moskvin et al., 2004; Lu and Stemler, 2002) have been insufficiently precise; further, this dehydration activity has been even much lower (∼0.02 - 0.07 and ∼0.26 W-A un.·mg^-1^ of Chl, respectively) than the hydration activity. Nevertheless, it was shown that the hydration activity was significant for providing maximal O_2_ evolution activity of PSII, isolated not only from higher plants but also from the green alga *Chlamydomonas reinhardtii*, which has its own CA - CAH3 associated with its PSII (Shitov et al., 2018, 2011; Terentyev et al., 2019). However, we are of the opinion that it is necessary to use PSII preparations, with high electron transport activity, to ensure high CA activity. Thus, PSII preparations (preps) must be used in the pH range of 6.0 – 6.5, which is known to be optimal for its O_2_ evolution (Schiller and Dau, 2000). Unfortunately, all the previous CA activity measurements had been carried out either at too high or too low pHs than needed, i.e., in the pH 8.0 – 8.3 range (the hydration (Shitov et al., 2018; Lu and Stemler, 2002; Khristin et al., 2004; Ignatova et al., 2006; Rudenko et al., 2007; Ignatova et al., 2011; Fedorchuk et al., 2014; Shitov et al., 2009, 2011)), or at pH 7.5 (the hydration and the dehydration (McConnell et al., 2007)), or both at pH 5.5 (Lu and Stemler, 2002) and at рН 6.8 (Moskvin et al., 2004) (the dehydration). Thus, to test the above hypothesis, a new, accurate, reproducible approach must be used for measurements of PSII CA activity at pH 6.0 – 6.5.

In 1997, Shingles and Moroney (Shingles and Moroney, 1997) indeed measured the dehydration activity of the most active α-CAs at pH 6.0, by using a highly sensitive fluorescent pH-indicator pyranine (8-hydroxy-pyrene-1,3,6-trisulfonate) in concert with a stop-flow technique. They found that CA preparations did not change the intensity of pyranine fluorescence, showing that the dye had no inhibitory effect on the activity of CAs (Shingles and Moroney, 1997). Thus, Shingles and Moroney concluded that the use of pyranine at pH 6.0 was a sensitive and accurate method for measuring high rates of HCO_3_^-^ dehydration for very active CAs. Below we describe how we have optimized the approach of Shingles and Moroney (Shingles and Moroney, 1997) for the measurements of HCO_3_^-^ dehydration at pH 6.5 in PSII from higher plants.

In this paper, we have answered two principal questions: have PSII from higher plants the true enzymatic CA activity with high efficiency and how connected photosynthetic and CA activities in PSII? We have proved the presence of highly efficient CA activity inside the PSII from *Pisum sativum*, showing that the dehydration activity at pH 6.5 is extremely high (∼650-3,550 times) than that measured at other pHs. Further, for the first time, we have determined catalytic properties of this activity: the maximal velocity (V_max_), the Michaelis constant (K_m_), the turnover number (k_cat_) and the specificity constant (k_cat_/K_m_); which, actually, showed high efficiency of HCO_3_^-^ dehydration in PSII that is comparable with the most active membrane bound α-CAs. More importantly, we have demonstrated here, for the first time, that the dehydration CA activity in PSII has, indeed, an enzymatic nature. Finally, we have found that CA activity interconnects with photosynthetic activity on the electron donor side of PSII. We propose that this interconnection realizes via the accelerating of removal of protons from the water-oxidizing complex (WOC) of PSII.

## 2. Results

The significance of CA activity for PSII and an enzymatic nature of this activity has been under debate for a very long time. McConnell et al. (McConnell et al., 2007) and Bricker and Frankel (Bricker and Frankel, 2011) presented the following arguments against an enzymatic nature of CA activity in PSII: (i) there is an absence of relationship between CA and photosynthetic activities in PSII (McConnell et al., 2007); (**ii**) there is contamination by water-soluble CAs from the stroma and/or from the cytoplasm (Bricker and Frankel, 2011); and **(iii**) a very low CA activity is observed in PSII (Bricker and Frankel, 2011). However, later in several papers (Shitov et al., 2011; Karacan et al., 2016; Rodionova et al., 2017; Shitov et al., 2018), we have shown clear relationship between CA activity, O_2_ evolution and photosynthetic electron transfer in PSII. In view of the earlier concerns (Bricker and Frankel, 2011; McConnell et al., 2007), we first checked the possibility of contamination of PSII (isolated by the procedure used in (Schiller and Dau, 2000), which is usually the one we use in our research) by water-soluble and membrane bound CAs. Second, we tested if PSII may have high CA activity at pH 6.5 (i.e. under the condition that is optimal for photochemical activity of this complex (Schiller and Dau, 2000)). Third, we have investigated here the kinetic parameters of this activity and its nature.

### 2.1. The test of contamination of PSII by other CAs

There are two possible sources of contamination of PSII by other CAs. **(1)** water-soluble CAs from the stroma of chloroplasts and from the lumen of thylakoids (in lesser degree, water-soluble CAs from cytoplasm of the cells) could “adhere” to PSII preps. **(2)** CA, associated with PSI (Rudenko et al., 2015), could contribute in CA activity of PSII preps (since most PSII preps usually contain small amounts of PSI).

#### On the contamination by water soluble CAs

The main reason of the contamination of thylakoid membranes (which is usually used to isolate PSII preps (Berthold et al., 1981; Schiller and Dau, 2000)) by water-soluble stromal CAs is the electrostatic and/or hydrophobic interactions between them. Moskvin (Moskvin et al., 2004) has demonstrated that these interactions are significantly impaired by repeated washing of thylakoid membranes (4 or more times), helping to obtain preps free of stromal water-soluble CAs. In current work, we isolated thylakoid membrane fragments enriched in PSII (BBY (Berthold-Babcock-Yocum)-particles (Berthold et al., 1981)) using 5 washings, which is expected to remove most of the water-soluble CAs. In addition, the detergent Triton X-100 (which is usually used (Berthold et al., 1981; Schiller and Dau, 2000) to prepare BBY-particles) is able to disrupt all the hydrophobic interactions between the BBY-particles and the water-soluble CAs. In general, the wide difference in the same activity of different BBYs, obtained by the same method, should indicate the contamination of samples by other CAs.

We observed only small differences in the CO_2_ hydration activities (Table 1, line 1; Wilbur-Aanderson units (mg Chl)^-1^ ((W-A un.) (mg Chl)^-1^)) among three BBY preps obtained, but this was simply due to the slightly different concentrations of photochemical reaction center II (RCII) in the used BBYs. However, when we expressed the activities in W-A un. related to the concentration of RCs (Table 1, line 2), two preps out of three showed no significant difference (according to T-Test, p = 0.16); these results are comparable with data expressed in the same units by Shitov et al. (Shitov et al., 2009) and are close to those obtained previously on samples from pea plants, obtained by several other authors (Shitov et al., 2018; Ignatova et al., 2006; Rudenko et al., 2007; Shitov et al., 2011). These data clearly show the low probability of contamination of our isolated BBYs by water soluble CAs.

**Table 1.**
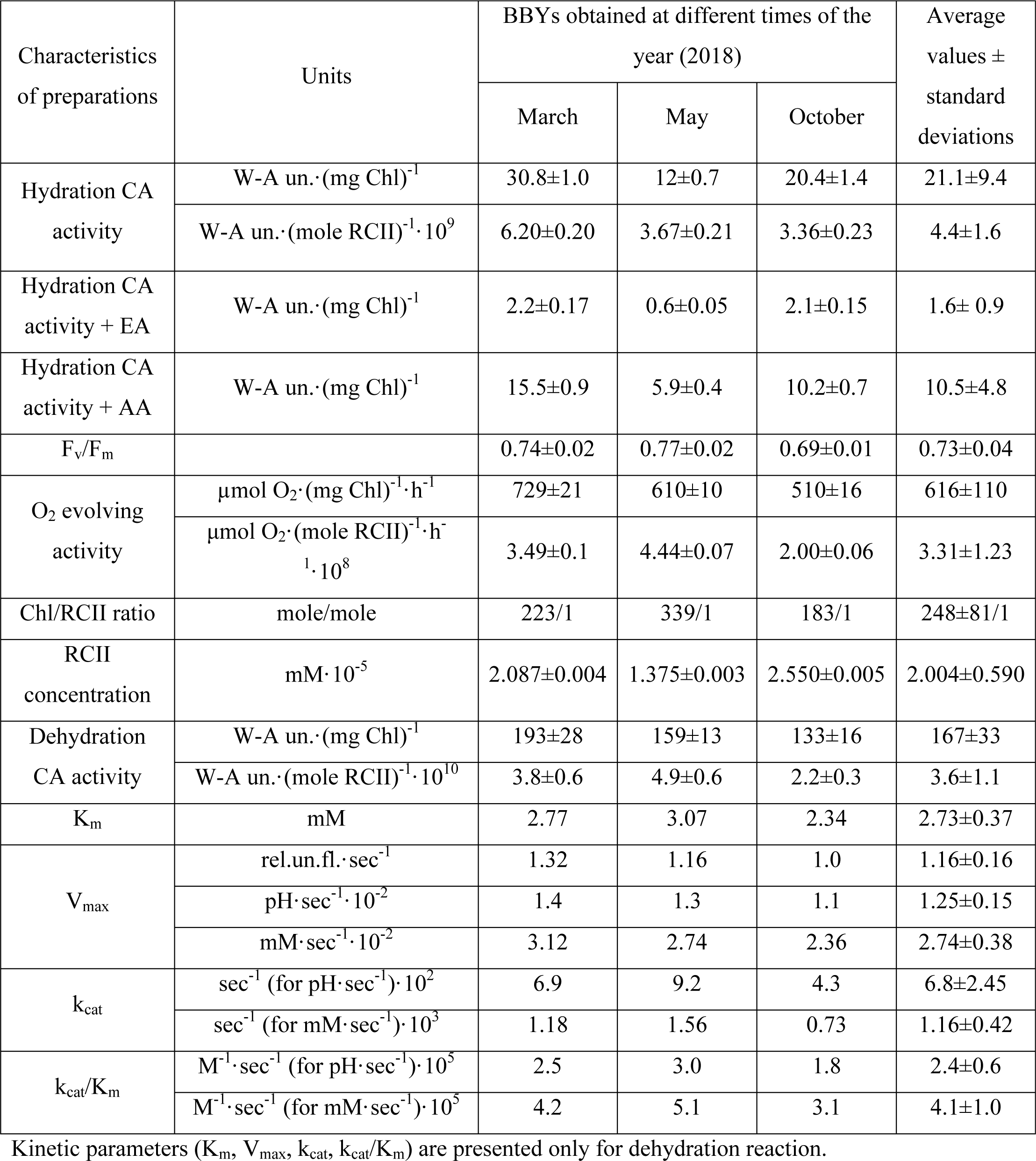
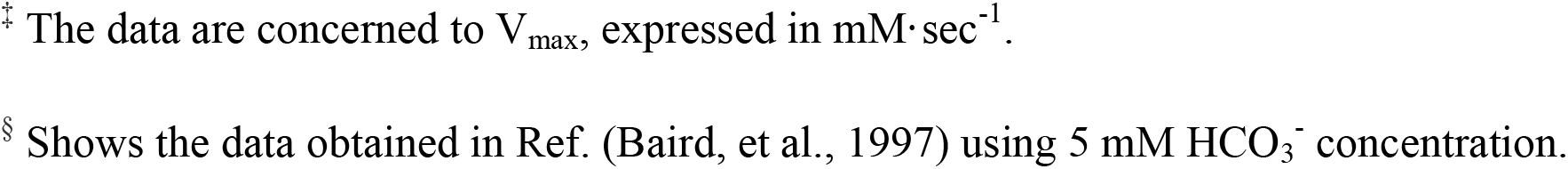
CA and photochemical activity of BBYs obtained at different times of the year.

#### On the contamination by CAs of PSI

Taking into account some heterogeneity in the properties of preps, further we tested the purity of the BBYs using the complex of 3 approaches (supplementing each other): (***a***) the test of CO_2_ hydration activity (which is described above); (***b***) the test of photochemical activity; and, (***c***) the influence of the well-known sulfonamide inhibitors (Supuran, 2008), such as ethoxyzolamide (EA) and acetazolamide (AA), on CO_2_ hydration activity. For this, we used 3 BBY preps, isolated from pea plants (*Pisum sativum*), grown at different times of the year (in March, May, and October 2018). Table 1 shows our results.

**(b)** A strong contamination of BBYs by CA associated with PSI may take place if and when the detergents used leave significant amount of PSI in the preparation. In this case, PSII preparation would show lower PSII photochemical activities (per chlorophyll (Chl)), because of the presence of Chls from PSI. Table 1 shows that the maximal quantum yield of our BBYs (inferred from F_v_/F_m_, the ratio of variable to maximum Chl fluorescence) was quite high (0.69 – 0.77), as well as the rate of oxygen evolution, ranged from 510 to 729 µmol O_2_·(mg Chl)^-1^·h^-1^, was also quite high. These results are characteristic of high photochemical activity (Berthold et al., 1981) of BBYs and this means that our preparations had only minor contamination of PSI, which could not significantly affect the measured CA activity, assigned to PSII.

**(*c*)** The use of sulfonamides is an effective approach to test the purity of PSII preps, since it is well known that CO_2_ hydration activity of PSII preps (obtained by different methods) has a unique response to the addition of acetazolamide (AA) and ethoxzolamide (EA). It is known that 10 µm EA inhibits CA activity of PSII almost completely, while 10 µm AA suppresses it only by 50% (Ignatova et al., 2006; Rudenko et al., 2007; Shitov et al., 2009, 2011, 2018). In contrast, the CA activity of PSI preps is known to be completely inhibited by 10 µm EA and by 10 µm AA (Ignatova et al., 2006; Rudenko et al., 2007). If PSII preps have any contamination by PSI, then the hydration CA activity of such preps should be inhibited by 10 µm AA more than 50%. This is also true if our PSIIs were contaminated with water soluble α-CAs and β-CA from the stroma (the most likely candidates for contamination of BBYs), since the CA activity of these CAs is also always suppressed by 10 µm EA and by 10 µm AA completely (Terentyev et al., 2019; Supuran, 2008; Rowlett, 2010). Our results, however, show that 10 µМ EA lowered CO_2_ hydration by 90-95%, while 10 µМ AA inhibited the activity only by 50% in all the BBYs, used here (Table 1, lines 3 and 4, respectively). These results indicate the unique response of the CA activity of our PSII preps to sulfonamides, which is in agreement with those published earlier (Ignatova et al., 2006; Rudenko et al., 2007; Shitov et al., 2009, 2011, 2018). Thus, the measured close amount of CO_2_ hydration activities, the identical inhibition of these activities by EA and AA (which is characteristic only for PSII) and the high photochemical activity of preparations, observed for all the BBYs used in our work, reliably demonstrate that the samples, used here, are free from any contamination by other CAs.

### 2.2. The investigation of CA activity of PSII at pH 6.5

To test our hypothesis that pH 6.5 is indeed the appropriate pH at which PSII would have high CA activity, we need a foolproof method for the measurement of CA activity at this pH in our PSII preps. Shingles and Moroney (Shingles and Moroney, 1997) had developed a very sensitive approach only for pH 6.0 and only for α-CAs. To investigate the CA activity of PSII at pH 6.5, we did the following: **(i)** modified the method, described by Shingles and Moroney, for pH 6.5; **(ii)** tested its sensitivity, using very active CA (whose catalytic properties are well known); (**iii**) measured CA activity of BBYs by our modified method; and (**iv**) determined the minimal concentration of PSII, which gave stable results. We describe below, step by step, details of the above.

#### 2.2.1. Adoption of the method described by Shingles and Moroney to pH 6.5

The maximal sensitivity of the measuring systems, using dyes (pH indicators), is usually characterized by the highest possible signal to noise ratio for absorption or fluorescence of the dye, which depends on the concentrations of the buffer and of the dye used (Kernohan, 1965; Pocker and Bjorkquist, 1977a; Shingles and Moroney, 1997; Nielsen and Fago, 2015). Data published in Refs (Kernohan, 1965; Pocker and Bjorkquist, 1977a; Shingles and Moroney, 1997; Nielsen and Fago, 2015) show that the concentration of the substrate determines the concentration of the buffer. Further, it is well known that the rate of a reaction is directly proportional to the substrate concentration only at low concentrations of the substrate (Lineweaver and Burk, 1934). We note that Shingles and Moroney (Shingles and Moroney, 1997) used the substrate up to 10 mM and the buffer (HEPES) at concentration 0.5 mM. In order to adapt their method to pH 6.5, we tested the concentration of the buffer (HEPES), from 2.5 to 10 mM; and of pyranine, from 1.8 to 11 µM, using spontaneous reaction of HCO_3_^-^ dehydration. We found that concentration 5 mM HEPES and 5.5 µM pyranine gave the highest signal to noise ratio (i.e. high sensitivity) for our measuring system. Then, we tested the concentrations of the substrate (HCO_3_^-^) from 1 to 10 mM. We obtained accurate reproducible results using 1.5 to 6 mM of HCO_3_^-^, which is close to that obtained by Shingles and Moroney (Shingles and Moroney, 1997)). Therefore, the low concentrations of the substrate (HCO_3_^-^) and of the buffer, obtained during the adoption of the published method, allowed us to accurately and reproducibly measure the spontaneous HCO_3_^-^ dehydration at pH 6.5.

#### 2.2.2. The test of the sensitivity of adapted method by α-CAII from bovine erythrocytes

α-CAII (carbonate hydrolase, EC 4.2.1.1, sequence accession number P00921, NCBI ID 9913) from bovine erythrocytes, being one of the most active CAs (in certain conditions), is a good enzyme for the test of the sensitivity of the method used in our work, since this CA was also used earlier by Shingles and Moroney (Shingles and Moroney, 1997), and its catalytic properties are well described by others (Kernohan, 1964; Pocker and Bjorkquist, 1977a). Our results show that the rate of HCO_3_^-^ dehydration for CAII is 20,192 ± 830 W-A un.·(mg protein)^-1^ or 5.87 (± 0.24)·10^11^ W-A un.·(mole protein)^-1^ (i.e. reaction center of the enzyme), which is very high, as compared to that published, e.g., by Karacan (Karacan et al., 2016). We examined the sensitivity of our measuring system by using different concentrations of the CAII (0.5, 0.25 and 0.125 µg·mL^-1^ (i.e., 1.725·10^-8^, 0.86·10^-8^ and 0.43·10^-8^ M)), and obtained accurate and reliable results even at 0.125 µg·mL^-1^ of the CAII (Supporting Information (SI) Fig S2). This low amount of CA is essentially the same to that reported by Shingles and Moroney (Shingles and Moroney, 1997), and it implies that our adapted method has a high sensitivity at pH 6.5 (which is as good as that described in Ref. (Shingles and Moroney, 1997)).

Using the measured initial rates of CAII activity at different HCO_3_^-^ concentrations, we derived the kinetic parameters (K_m_ and V_max_), and calculated k_cat_ as well as k_cat_/K_m_ (see Table 2, column 2 and SI section 1 (Figs S1–S3)). The K_m_ value (25.7 mM) for CAII, obtained here, is close to that published (Kernohan, 1964; Pocker and Bjorkquist, 1977a). The V_max_ for CAII is equal to 0.28 pH·sec^-1^, which is similar to that published by Shingles and Moroney (Shingles and Moroney, 1997). Based on the similarity of the obtained data (Table 2, column 2) to the results, described by others (Table 2, columns 3-7), and on the high sensitivity of the adapted method, we conclude that the method, described here, is certainly applicable for accurate measurements of high rates of HCO_3_^-^ dehydration reaction at pH 6.5.

**Table 2.**
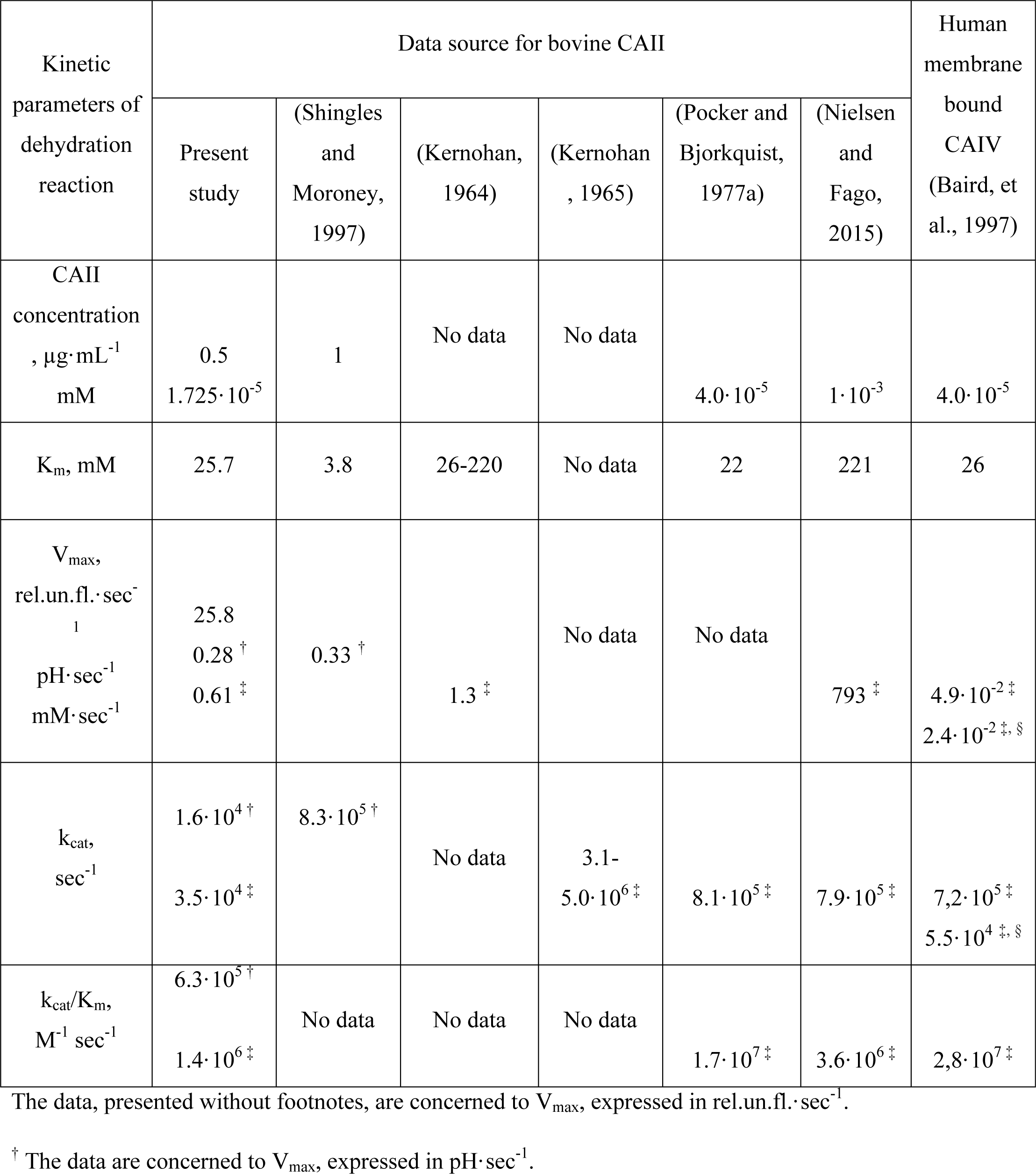
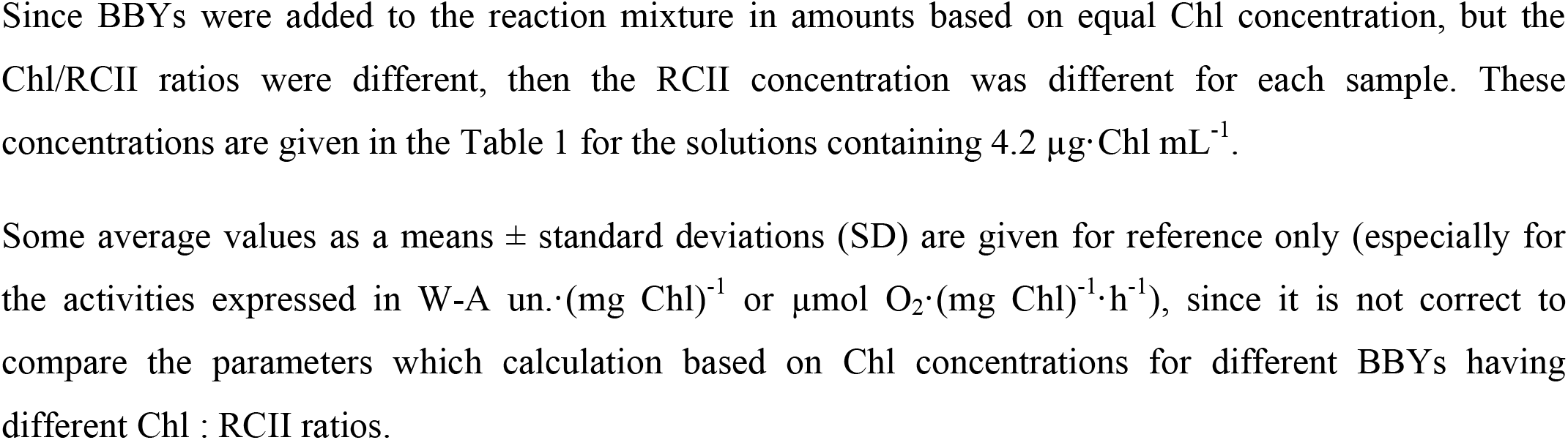
The comparison of bovine CAII kinetic parameters, obtained in the present work with those published previously.

| Kinetic parameters of dehydration reaction | Data source for bovine CAII |  |  |  |  |  | Human membrane bound CAIV (Baird, et al., 1997) |
| --- | --- | --- | --- | --- | --- | --- | --- |
|  | Present study | (Shingles and Moroney, 1997) | (Kernohan, 1964) | (Kernohan, 1965) | (Pocker and Bjorkquist, 1977a) | (Nielsen and Fago, 2015) |  |
| CAII concentration, $\mu\text{g}\cdot\text{mL}^{-1}$<br>mM | 0.5<br>$1.725\cdot 10^{-5}$ | 1 | No data | No data | $4.0\cdot 10^{-5}$ | $1\cdot 10^{-3}$ | $4.0\cdot 10^{-5}$ |
| $K_m$, mM | 25.7 | 3.8 | 26-220 | No data | 22 | 221 | 26 |
| $V_{\max}$,<br>$\text{rel.un.fl.}\cdot\text{sec}^{-1}$<br>$\text{pH}\cdot\text{sec}^{-1}$<br>$\text{mM}\cdot\text{sec}^{-1}$ | 25.8<br>$0.28^{\dagger}$<br>$0.61^{\ddagger}$ | $0.33^{\dagger}$ | $1.3^{\ddagger}$ | No data | No data | $793^{\ddagger}$ | $4.9\cdot 10^{-2\ddagger}$<br>$2.4\cdot 10^{-2\ddagger, \$}$ |
| $k_{\text{cat}}$,<br>$\text{sec}^{-1}$ | $1.6\cdot 10^4{}^{\dagger}$<br>$3.5\cdot 10^4{}^{\ddagger}$ | $8.3\cdot 10^5{}^{\dagger}$ | No data | 3.1-<br>$5.0\cdot 10^6{}^{\ddagger}$ | $8.1\cdot 10^5{}^{\ddagger}$ | $7.9\cdot 10^5{}^{\ddagger}$ | $7.2\cdot 10^5{}^{\ddagger}$<br>$5.5\cdot 10^4{}^{\ddagger, \$}$ |
| $k_{\text{cat}}/K_m$,<br>$\text{M}^{-1}\text{sec}^{-1}$ | $6.3\cdot 10^5{}^{\dagger}$<br>$1.4\cdot 10^6{}^{\ddagger}$ | No data | No data | No data | $1.7\cdot 10^7{}^{\ddagger}$ | $3.6\cdot 10^6{}^{\ddagger}$ | $2.8\cdot 10^7{}^{\ddagger}$ |

The data, presented without footnotes, are concerned to $V_{\max}$ , expressed in $\text{rel.un.fl.}\cdot\text{sec}^{-1}$ .
$^{\dagger}$ The data are concerned to $V_{\max}$ , expressed in $\text{pH}\cdot\text{sec}^{-1}$ .

#### 2.2.3. The investigation of HCO_3_^-^ dehydration activity of BBYs, using the method adapted for measurements at pH 6.5

The dehydration activity of BBYs was found to be 167 ± 33 W-A un.·(mg Chl)^-1^ (Table 1, line 10). This activity is ∼650 and ∼3,550 times higher than the dehydration activity measured at pH 5.5 (Lu and Stemler, 2002) and at pH 6.7 (Moskvin et al., 2004), respectively, and ∼8 times higher than the hydration activity, measured at pH 8.3 (Table 1, line 1). Thus, the CA activity of PSII at pH 6.5, shown here, is greater than at lower and at higher pHs, which is in agreement with our proposal (see “Introduction”).

To compare HCO_3_^-^ dehydration activity of BBYs with that of CAII (Fig. 1) we used equal concentrations of active centers of CA and BBYs. Since we used the concentration of active centers of CAII equal to 1.725·10^-8^ M (0.5 µg·mL^-1^), as was described in Ref. (Shingles and Moroney, 1997); in most experiments, we used the concentration of BBYs which was equal to the concentration of RCII ∼2·10^-8^ M (corresponding to 4.2 (µg Chl)·mL^-1^). The activity of CA in BBYs ((3.6 ± 1.1)·10^10^ W-A un.·(mole RCII)^-1^ (Table 1, line 11)) was found to be ∼ 16 times lower than that of CAII (Table 2, column 2). This result is not an indication of the low CA activity of PSII, in principle, because there are membrane-bound α-CAs (for example, CAXV from mouse (Hilvo et al., 2005)), that have ∼30 times lower CA activity. Thus, here we show (using the method of measurements that is new for PSII), for the first time, that the CA activity of PSII at pH 6.5 is significantly higher than that of some α-CAs, which implies, at least, the enzymatic nature of catalysis of HCO_3_^-^ dehydration in PSII. Below, in section 2.3., we will present the additional evidence of enzymatic nature of CA activity in PSII.

**Figure 1.**
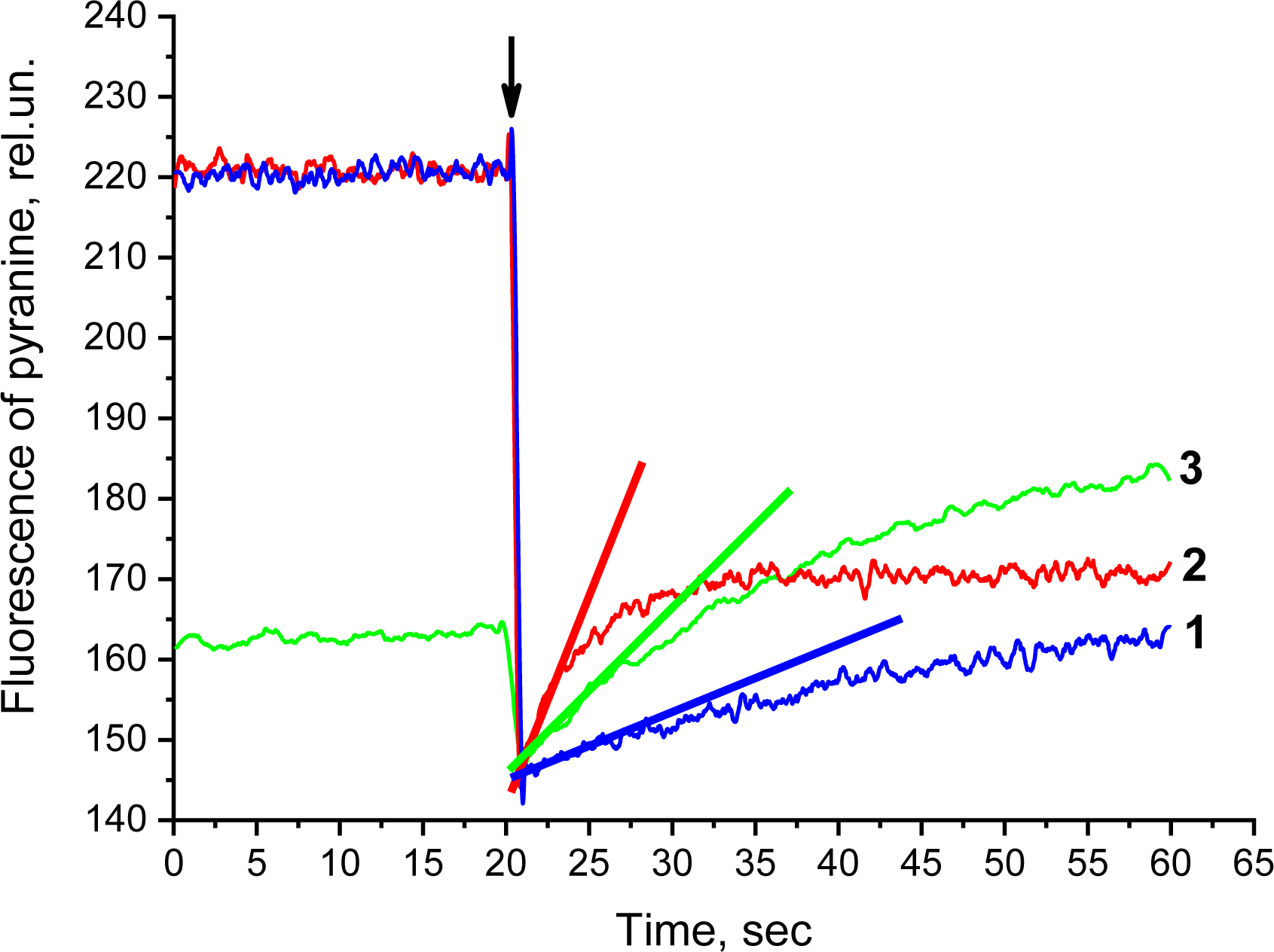
Kinetics of pyranine fluorescence changes related to the HCO_3_^-^ dehydration activity of BBY-particles and CAII. For these experiments, we used BBY preparation obtained in March 2018. The curve 1 shows the pyranine fluorescence changes in the buffer in the absence of biological samples (spontaneous reaction). The curve 2 shows pyranine fluorescence changes in the presence of bovine CAII (CAII catalyzed reaction) at final concentration of the protein equal to 0.5 µg·mL^-1^ (1.725·10^-8^ M). The curve 3 displays the pyranine fluorescence changes in the presence of BBY (PSII catalyzed reaction) at final concentration of BBY containing 4.2 (µg Chl)·mL^-1^. Straight lines represent the initial rates of the reaction of HCO_3_^-^ dehydration. The arrow indicates the moment of the addition of HCO_3_^-^. The concentration of the substrate (HCO_3_^-^) was 1 mM. The curves were smoothed by algorithm of Savitzky-Golay 35 points.

#### 2.1.4. The test of the sensitivity of our adapted method for PSII

We noted that the presence of BBY decreased the initial level of pyranine fluorescence, before the addition of HCO_3_^-^ (Fig. 1, curve 3). The decrease of pyranine fluorescence was found to be dependent on the concentration of BBY (Figs 2 and 3). Before checking the sensitivity of the method for PSII, we should **(1)** determine the reason of the decrease of the fluorescence of pyranine and **(2)** test whether this decrease affects the CA activity.

**Figure 2.**
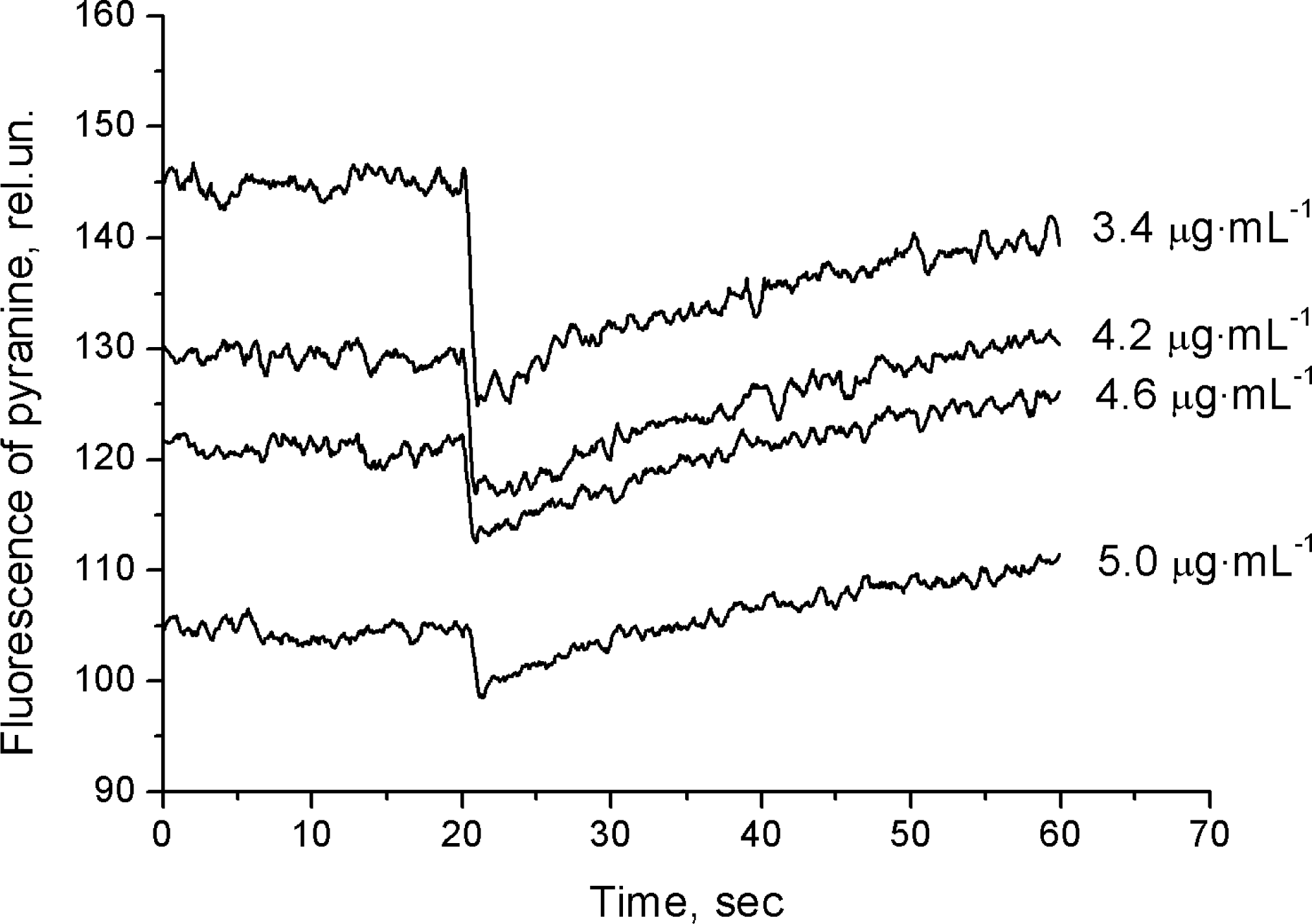
Kinetics of pyranine fluorescence changes related to the HCO_3_^-^ dehydration activity of BBYs used at different concentrations (given in terms of [Chl]). The final Chl concentration in the reaction mixture are shown on the right side of the curves; they are given in terms of µg Chl·mL^-1^. The concentration of the substrate (HCO_3_^-^) was 4 mM. For these experiments BBY preparation, obtained in May 2018 was used.

**Figure 3.**
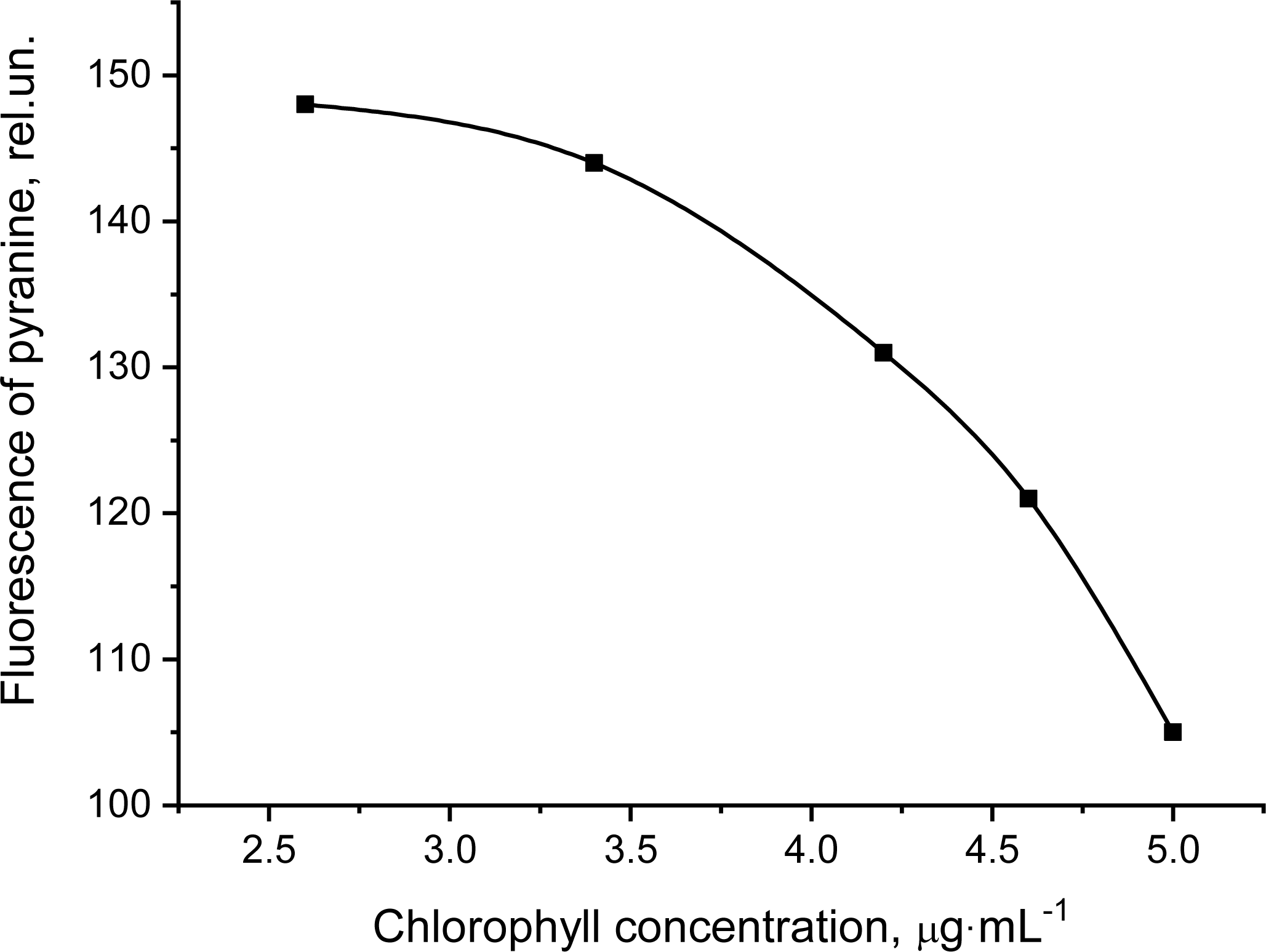
Pyranine fluorescence as a function of Chl concentration of the BBYs used; data show that pyranine fluorescence decreases with increasing concentration of BBY preparations. Pyranine fluorescence was measured before the addition of bicarbonate (i.e. before carbonic anhydrase reaction was started). The final concentrations of BBY were, in terms of Chl, 2.6, 3.4, 4.2, 4.6 and 5.0 (µg Chl)·mL^-1^. For these experiments, we used BBY preparation obtained in May 2018.

**(1)** We noted that the decrease of pyranine fluorescence was proportional to the concentration of Chl in the reaction mixture (Figs 2 and 3), therefore we propose that this decrease occurs due to the presence of photosynthetic pigments of PSII in the reaction mixture. Actually, we have found (Fig. 4) that photosynthetic pigments of PSII are able to absorb the light at the wavelength of pyranine excitation (466 nm), which may result in the decrease of pyranine fluorescence (for more information see SI). To test this hypothesis, we have extracted the pigments from BBY and further added obtained extract (at the same concentration of Chl as in BBY (4.2 (µg Chl)·mL^-1^)) in the chamber B (for more information see section 4.2. of “Materials and Methods”) instead of BBYs. When the extract was added, the pyranine fluorescence, before the addition of HCO_3_^-^, was found to be lowered (Fig. 5A, curve 2) similarly to that of the BBYs (Fig. 1, curve 3), which indicates that the decrease of pyranine fluorescence really caused by the presence of photosynthetic pigments. (For additional information on the influence of photosynthetic pigments of PSII on the fluorescence of pyranine, see SI sections 2 and 3 (Figs S4–S7)).

**Figure 4.**
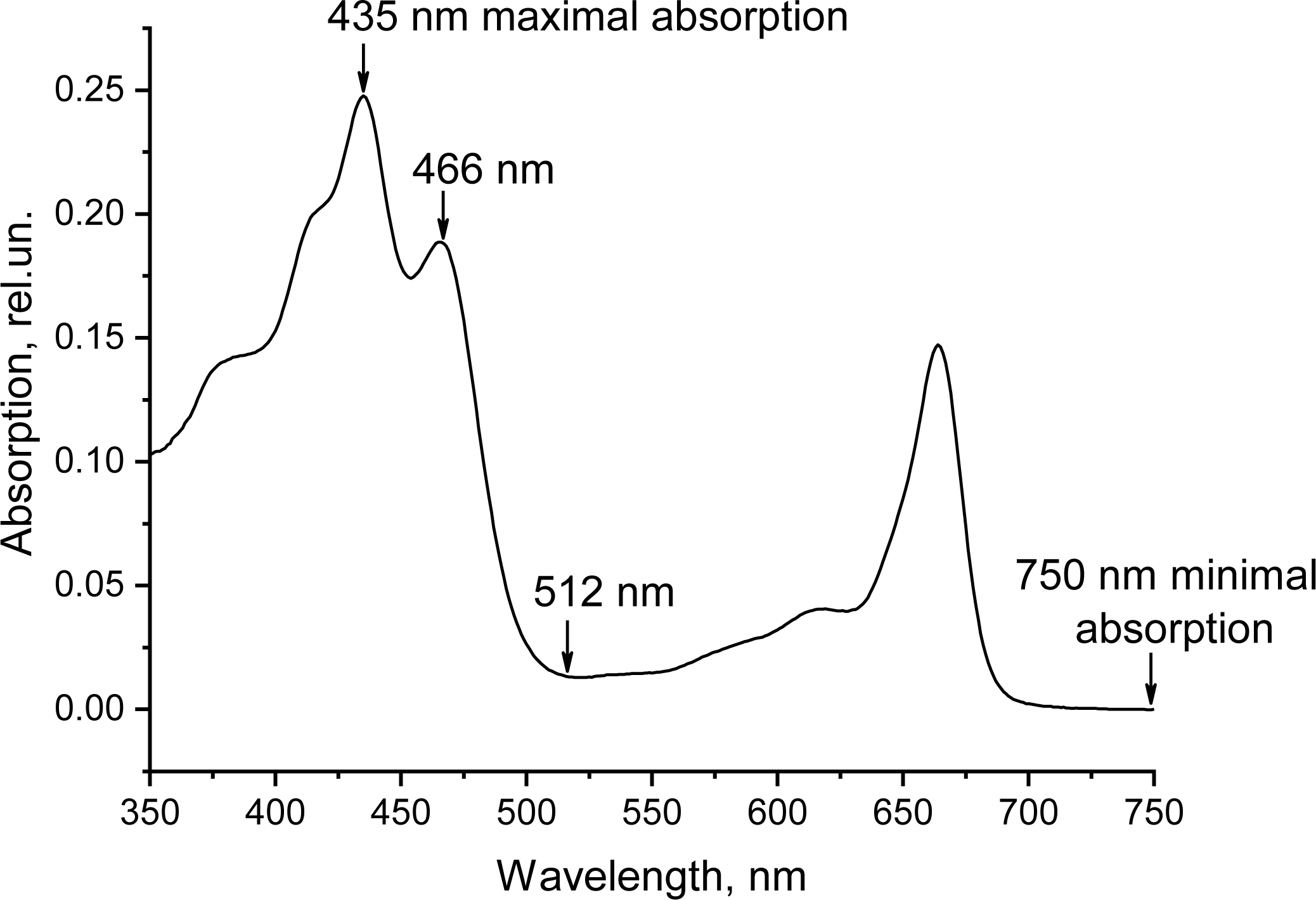
The absorption spectrum of photosynthetic pigments, obtained from BBY preps, and suspended in 96% ethanol. The absorption peak was at 435 nm, where the absorbance was 0.2475; at 512 nm (the emission of pyranine), this value was 0.0143 (which is 5.8% of the maximum); at 466 nm (the excitation of pyranine), this value was 0.1890 (which is 76% of the maximum). For these experiments, we used BBY preparation obtained in May 2018.

**Figure 5.**
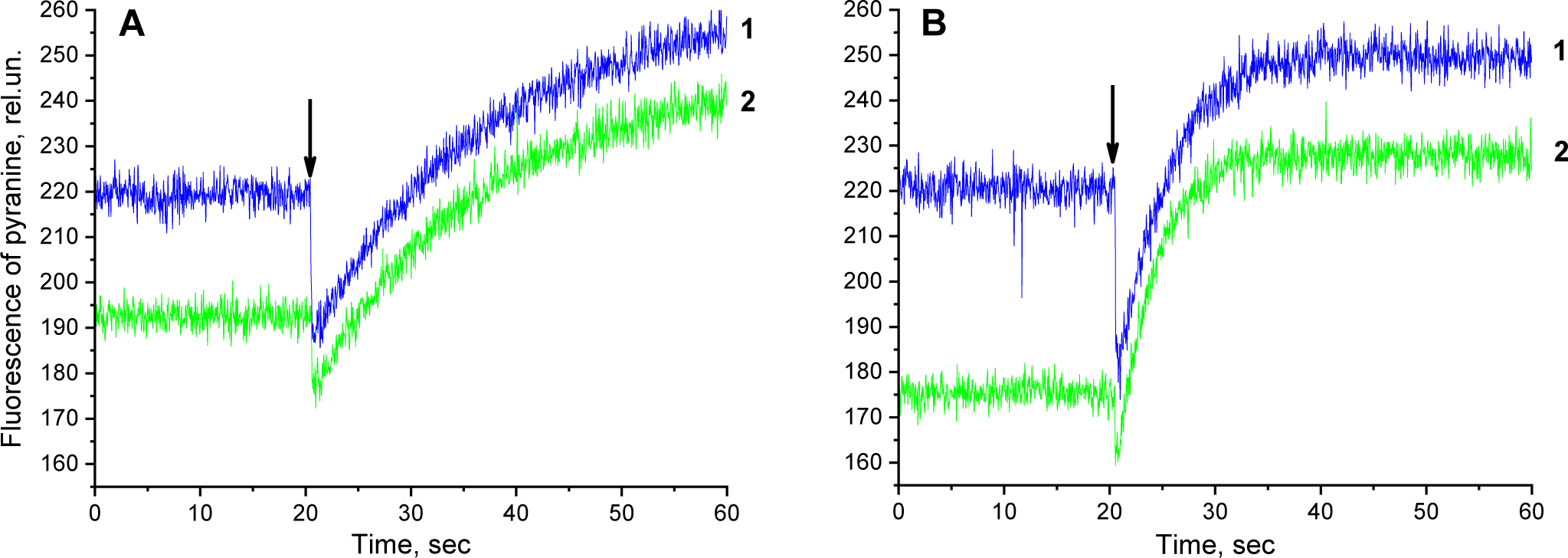
The influence of Chl extract on pyranine fluorescence changes, related to the HCO_3_^-^ dehydration. (A) the spontaneous reaction, (B) the reaction, catalyzed by CAII. Curve 1 shows the kinetics of pyranine fluorescence changes in 5 mM HEPES buffer with added ethanol (10%). Curve 2 presents the kinetics of pyranine fluorescence changes in 5 mM HEPES buffer with added Chl extract in ethanol (the concentration of ethanol in the mixture was also 10%). The arrow indicates the moment of the addition of HCO_3_^-^. The concentration of bicarbonate was 4 mM. The final concentration of CAII was 0.5 µg·mL^-1^. The final concentration of Chl was 4.2 µg·mL^-1^.

**(2)** To test the possible influence of photosynthetic pigments on the dehydration reaction, we used CAII from bovine erythrocytes, since we know exactly the value of the activity of this CA, and even a little change in the activity of such an active CA could be easily observed. For this experiment, we added the extract of pigments to the chamber B along with the addition of CAII. We noted that, in the case of the simultaneous presence of CAII and the extract of pigments (Fig. 5B, curve 2), the fluorescence of pyranine, before the addition of HCO_3_^-^, changed close to that of the spontaneous reaction (Fig. 5A, curve 2), which indicates that the presence of CAII has no effect on the fluorescence of pyranine if we add to it the extract of photosynthetic pigments. In addition, in this case (Fig. 5B, curve 2), the fluorescence of pyranine, before the addition of HCO_3_^-^, also changes similarly to that of the BBYs (Fig. 1, curve 3), which confirm our above suggestion that only photosynthetic pigments actually lowered the fluorescence of pyranine in the presence of BBYs. The presence of the pigments does not affect the initial rate and the amplitude of fluorescence changes of pyranine, for both the spontaneous and the enzymatic reactions, after the addition of HCO_3_^-^ to the reaction mixture (Figs 5A and 5B, respectively). Thus, we conclude that the presence of photosynthetic pigments of PSII does not influence the activity of CAII and, consequently these pigments does not affect the sensitivity of the method.

To test the sensitivity of the method, used here for PSII, we measured the activity of BBY at different Chl concentrations (ranging from 2.6 to 5.0 (µg Chl)·mL^-1^). The linear dependence of the reaction rate on such a low BBY concentrations (Fig. 6) shows the high sensitivity of this method for our PSII samples; we note that this sensitivity is ∼50 times higher for HCO_3_^-^ dehydration than that for CO_2_ hydration. Thus, the method is sensitive enough for measurements of HCO_3_^-^ dehydration activity in PSII preps.

**Figure 6.**
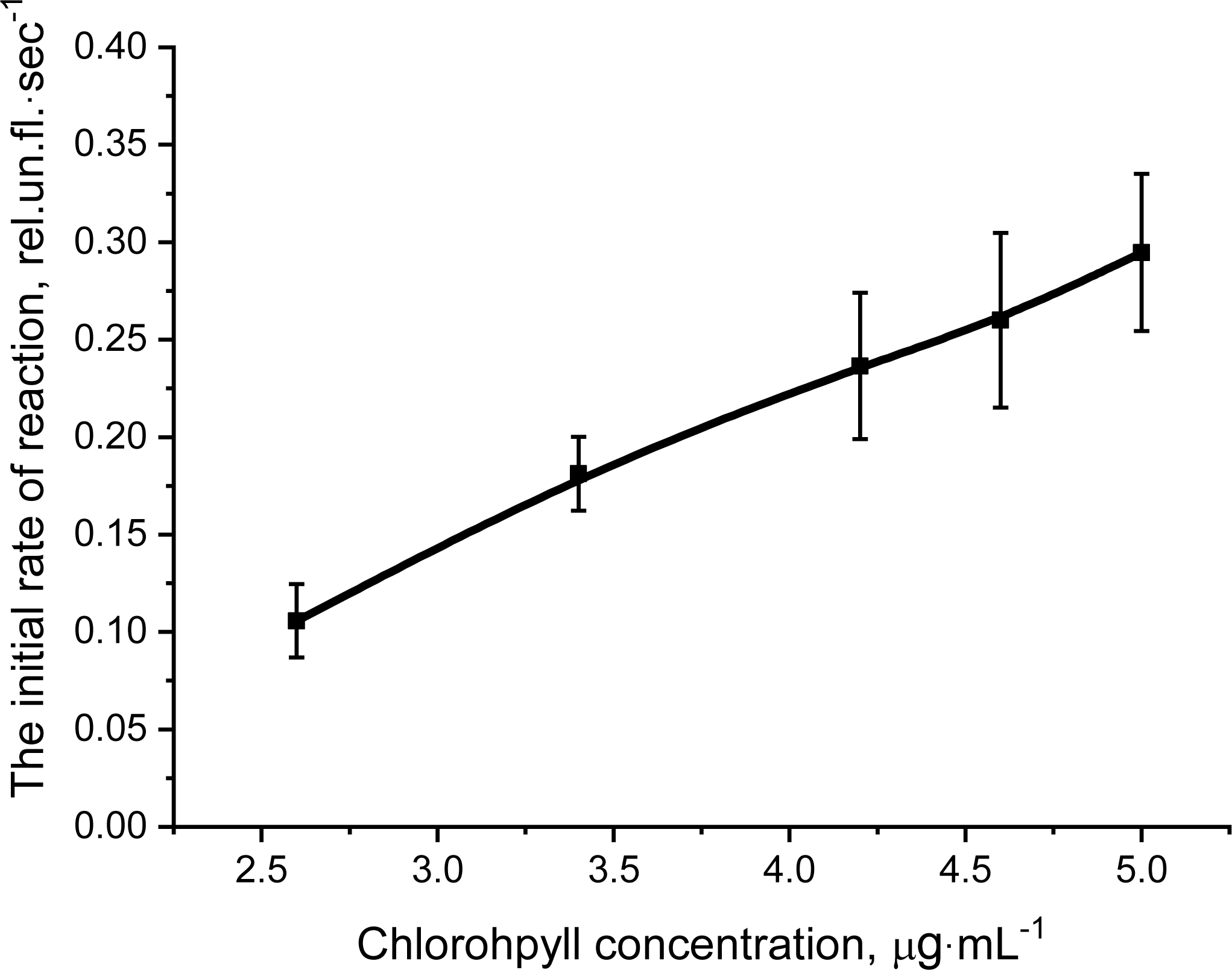
The dependence of initial rate of pyranine fluorescence changes on the amount of BBY, given in terms of their chlorophyll concentration. For these experiments we used BBY preps, obtained in March 2018, in May 2018 and in October 2018. The concentration of bicarbonate was 4 mM. The final concentrations of BBYs had 2.6, 3.4, 4.2, 4.6 and 5.0 (µg Chl)·mL^-1^. Each point is an average of at least 3 separate experiments with calculated standard deviation (SD).

### 2.3. The nature of HCO_3_^-^ dehydration and its kinetic parameters in PSII

To investigate the nature of the reaction, studied here, we should determine the concentration of the catalyst. We note that the HCO_3_^-^ dehydration activity of PSII (Table 1, line 11), expressed in W-A un.·(mole RCII)^-1^, is directly proportional to the number of Chl molecules bound to the RCII, i.e., the activity is proportional to the ratio of Chl/RCII (Table 1, line 8). Using the plot of “CA activity” vs. “the ratio of Chl/RCII” (Fig. 7), we note that ∼ 15 molecules of Chl is the minimal number of molecules per one active center of CA in our BBY preps. It is known (van Leeuwen et al., 1991) that the RCII of PSII (the minimal complex that is capable of photochemical reactions) contains six molecules of Chl, whereas the next largest complex, the Core-complex, contains 35 to 80 molecules of Chl. Since, the number of 15 molecules of Chl per one active center of CA is closest to that of Chl in RCII, we speculate that the one active center of CA in PSII corresponds to (or is related to) one photochemical RCII. For further analysis of our data, we will use the concentration of RCII as a parameter of [E_0_] in the calculations of k_cat_ and k_cat_/K_m_ for HCO_3_^-^ dehydration in PSII (see below).

**Figure 7.**
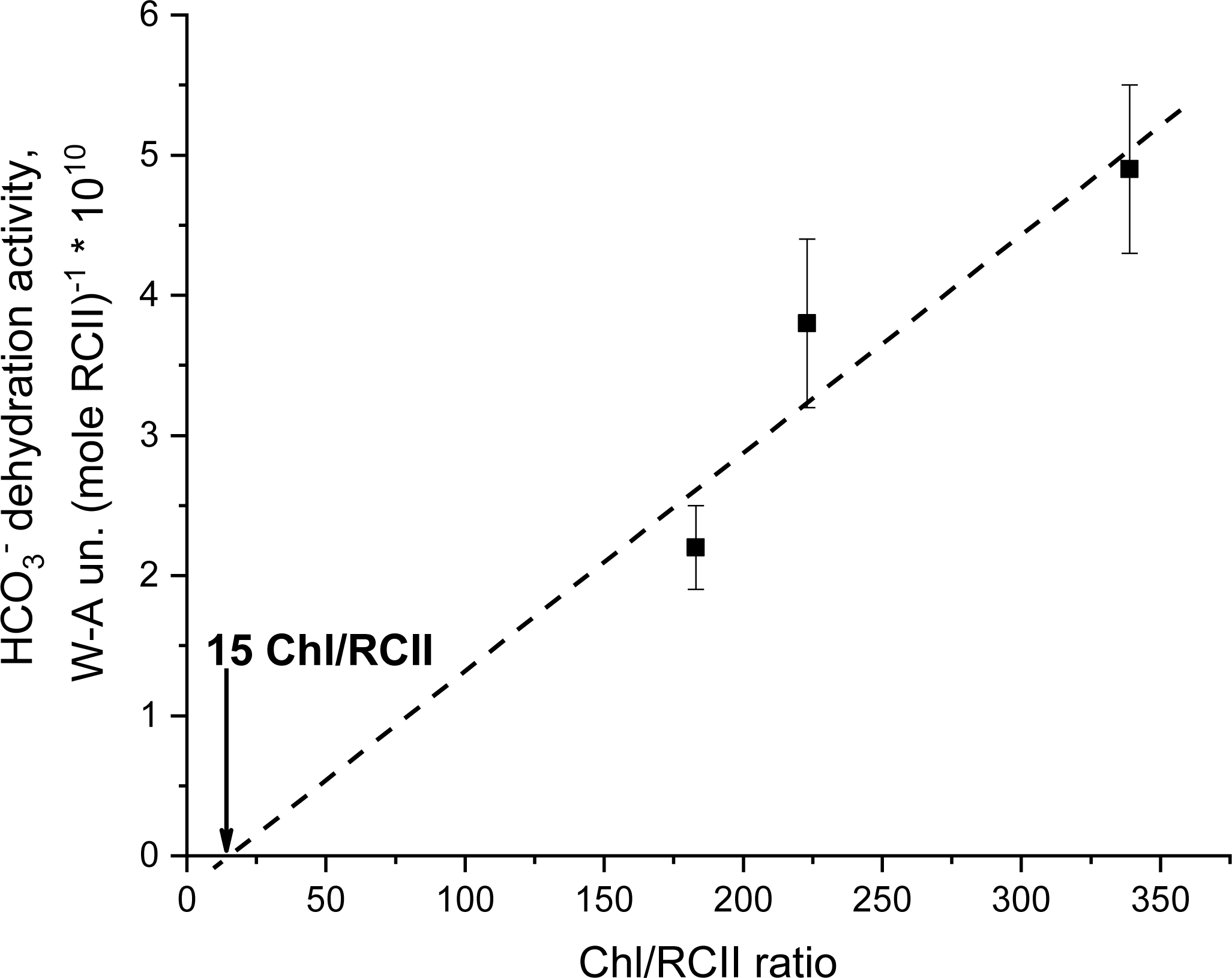
The dependence of HCO_3_^-^ dehydration activity of BBYs as a function of the ratio of Chl/RCII in these preps. For these experiments we used BBY preps, obtained in March 2018, in May 2018 and in October 2018. The final concentration of BBYs, given as Chl·mL^-1^, was 4.2 (µg Chl)·mL^-1^. The concentration of bicarbonate was 4 mM. Each point is an average of at least 3 separate experiments with calculated SD. Adj. R-Square of the line plotting was 0.72.

Fig. 1 shows that HCO_3_^-^ dehydration, in the presence of BBY, occurs faster than in the spontaneous reaction, which implies, at least, that BBY catalyzes this reaction. This catalysis may indicate the enzymatic nature of HCO_3_^-^ dehydration in PSII, but this suggestion need additional evidences. Below we present these evidences of enzymatic nature of CA activity in PSII.

We note that the kinetics of HCO_3_^-^ dehydration in PSII (Fig. 1, curve 3) is similar to that of CAII shown in Fig. 1, curve 2 and, especially, to that of CAII, presented by Shingles and Moroney (Shingles and Moroney, 1997), who had described this kinetics as “catalyzed reaction [which] follows first-order kinetics”. Since PSII also catalyzes HCO_3_^-^ dehydration (see above), following first-order kinetics, these facts imply that (1) the same first order reaction (Eq. 3) is involved in both CAII and in PSII; (2) the same mechanism of this reaction works in both cases; and (3) the same (enzymatic) nature of catalysis occurs in both cases. Thus, the observed kinetics of HCO_3_^-^ dehydration in BBYs implies the enzymatic nature of CA activity in PSII.

The following feature also indicate the enzymatic nature of HCO_3_^-^ dehydration in PSII. The plot of the “initial rate” vs. the “initial substrate concentration”, obtained for BBYs at seven substrate concentrations (Fig. 8), shows “saturation” at concentrations of HCO_3_^-^ greater than 4.5 mM, which is similar to that of some of the enzymes described by Michaelis and Menten (Johnson and Goody, 2011). As postulated by Michaelis and Menten (Johnson and Goody, 2011), the saturation of the rate is the characteristic of enzymatic reactions at increased substrate concentrations. This is right only if the concentration of the enzyme is significantly lower than the concentration of the substrate (Johnson and Goody, 2011), because in this case molecules of a substrate fully saturate all the active centers of an enzyme (Lineweaver and Burk, 1934), preventing further increase of the rate. In our experiments, the condition, postulated in Ref. (Johnson and Goody, 2011), is achieved, since the concentration of active centers of CA in BBYs (∼2·10^-8^ M) was extremely low (5 orders of magnitude) than saturating concentrations of the substrate (5 – 6 mM). Therefore, we conclude that HCO_3_^-^ dehydration (and CA activity, in general) has indeed an enzymatic nature in PSII.

**Figure 8.**
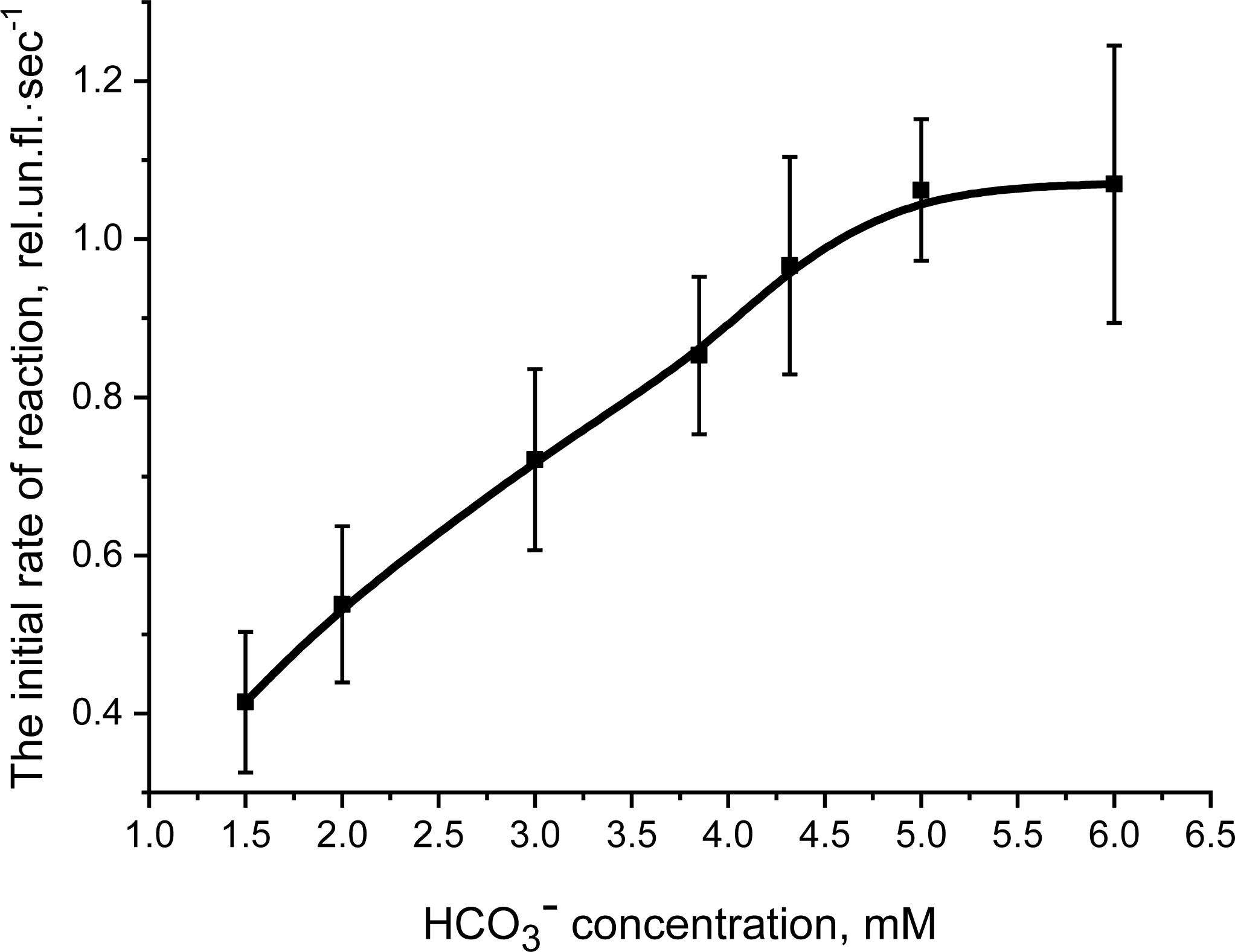
The dependence of initial rate of HCO_3_^-^ dehydration activity of PSII on the substrate (bicarbonate) concentration. For these experiments we used BBY preps, obtained in March 2018, in May 2018 and in October 2018. The concentrations of bicarbonate (for each BBY preparation) were 1.5, 2, 3, 3.85, 4.3, 5 and 6 mM. The final concentration of BBY, in units of Chl per mL, was 4.2 (µg Chl)·mL^-1^. Each point is an average of at least 3 separate experiments with calculated SD.

The main indicator of the enzymatic nature of the reaction is the observed high V_max_, k_cat_ and k_cat_/K_m_, which were shown to exceed the kinetic parameters of spontaneous reaction. We have calculated K_m_ and V_max_ for different BBYs using Lineweaver-Burk plots, that are shown in Fig. 9, obtained results we present in Table 1, in which we summarize all the kinetic parameters, obtained for BBY preparations. We found that V_max_ for BBYs, ranged from 2.4·10^-2^ to 3.1·10^-2^ mM·sec^-1^, which is close to that of membrane bound α-CAIV (especially at low (5 mM) HCO_3_^-^ concentration (Baird, et al., 1997) (Table 2, column 8)). We note that α-CAIV belongs to the most active transmembrane and membrane-bound α-CAs (Baird, et al., 1997; Supuran, 2008), and, thus, our data imply that the CA in PSII is also highly effective.

**Figure 9.**
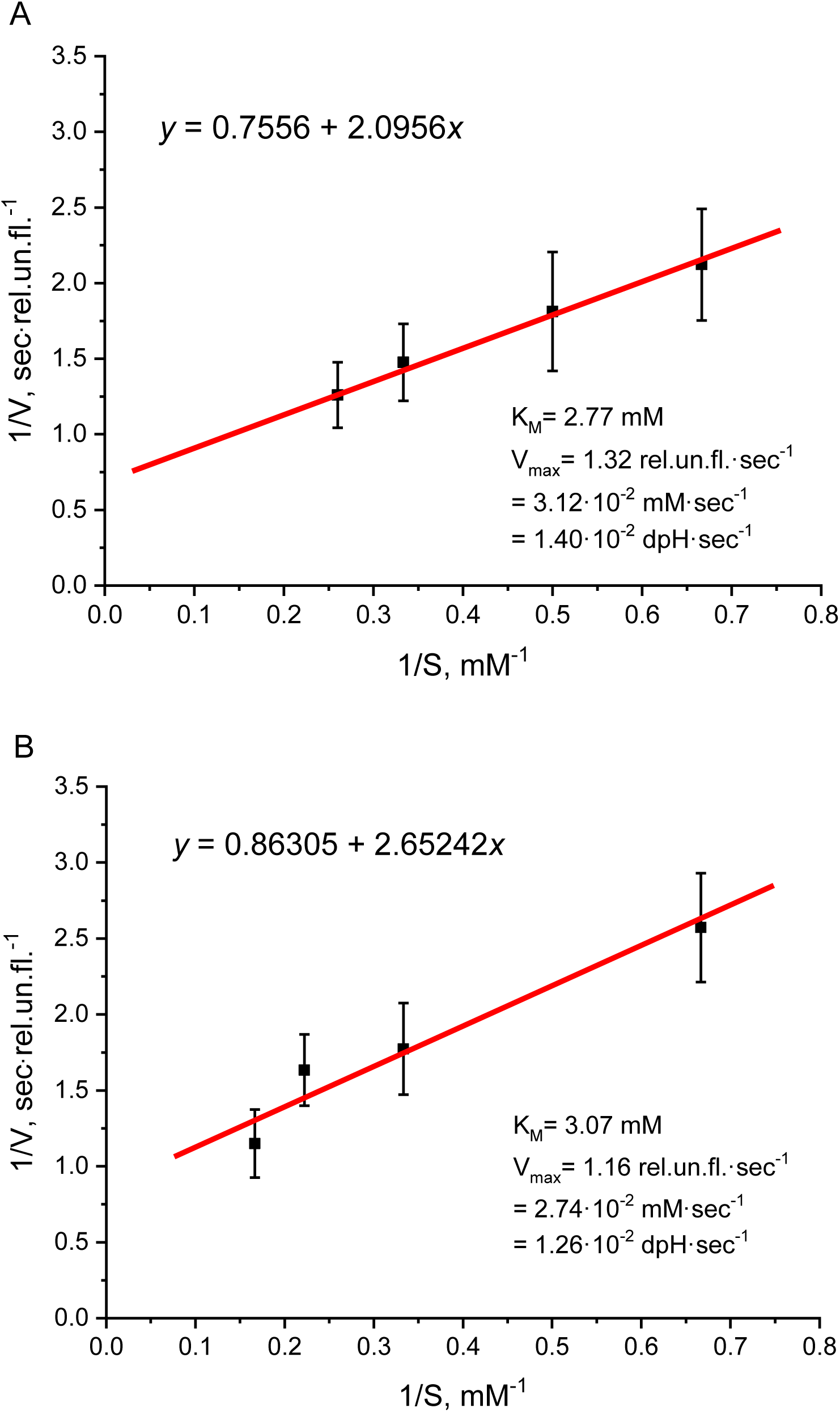
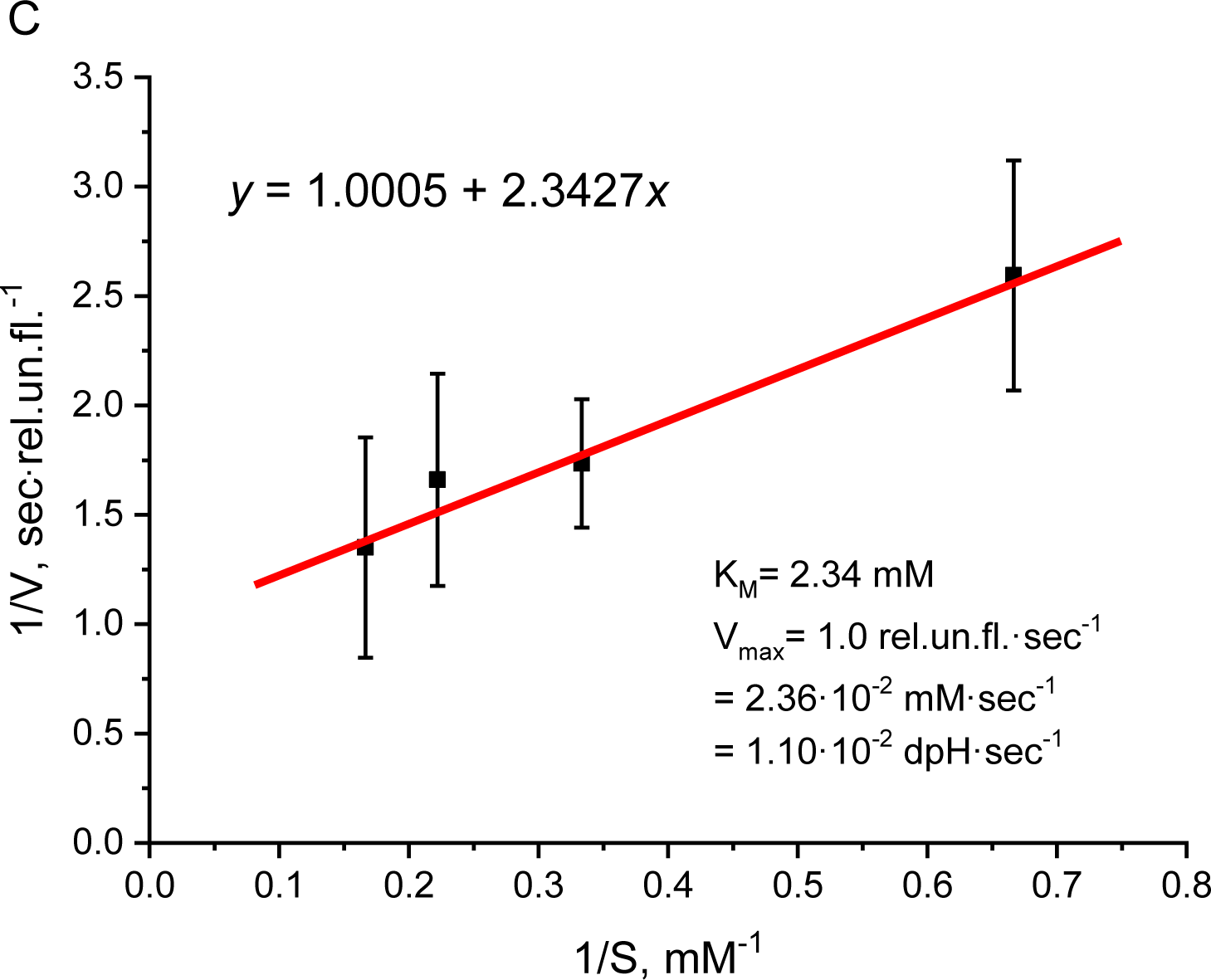
The Lineweaver and Burk plots for the initial rate of HCO_3_^-^ dehydration activity as a function of the substrate (HCO_3_^-^) concentration, obtained for BBY preparations isolated at different times of the year. (A) BBY preparation, obtained in March 2018, the concentrations of bicarbonate were 1.5, 2.0, 3.0 and 3.85 mM; (B) BBY preparation, obtained in May 2018, the concentrations of bicarbonate were 1.5, 3.0, 4.5 and 6.0 mM; (C) BBY preparation, obtained in October 2018, the concentrations of bicarbonate were 1.5, 3.0, 4.5 and 6.0 mM. We used the equations of the plots, shown in the above curves, to calculate K_m_ and V_max_, and these results are shown in Table 1. The final concentration of BBYs was 4.2 (µg Chl)·mL^-1^. Each point was an average of at least 3 separate experiments with calculated SD. Adj. R-Squares of line plotting were 0.95, 0.86 and 0.94, respectively.

Further, using K_m_ and V_max_ (presented in Table 1), we have calculated k_cat_ and k_cat_/K_m_ (the obtained results are also presented in Table 1). These results indicate that k_cat_ for HCO_3_^-^ dehydration in BBYs exceeds the rate constant for the spontaneous reaction (that equals 18 sec^-1^ (Pocker and Bjorkquist, 1977b)) by ∼67 times, which indicates that there is indeed catalysis of this reaction in PSII. CA activity of PSII, with k_cat_=1.2·10^3^ sec^-1^ and k_cat_/K_m_=4.1·10^5^ M^-1^ sec^-1^, is only an order of magnitude lower than that of the most active CAII. Moreover, we note that some CAs have considerably lesser efficiency of catalysis than in PSII; this is so for the water-soluble CAIII (Baird, et al., 1997; Supuran, 2008) that has k_cat_/K_m_ ∼50 fold lesser value than that of PSII. Thus, we conclude that the dehydration activity of PSII has an enzymatic nature with high efficiency of catalysis.

We note that PSIIs have low K_m_ (2.7 mM), which is almost 10 times lower than that of some α-CAs (Table 2) and at least 20 times lower than that of stromal β-CA (Johansson and Forsman, 1993). This finding implies that the BBYs, used in this work, were quite pure, i.e., they were not contaminated by α- and β-CAs. If BBYs, used here, were contaminated by α- and β-CAs, then these preparations would have had much higher K_m_ and, consequently, there would have been larger differences between them (at least, in some of them). This is not the case (i.e., the values of the K_m_ are very similar and small in all the BBYs (Table 1, line 12)), and, thus we confirm our conclusion that the BBYs, used in our research, were free of other CAs.

Thus, using very sensitive method of the measurements of CA activity (which is new for PSII), we have estimated (Table 1) that PSII, from the pea leaves, enzymatically, with high efficiency, accelerates HCO_3_^-^ dehydration at pH 6.5 (optimal for PSII photochemistry (Schiller and Dau, 2000)).

## 3 Discussion

The investigation of the function of an enzyme consists of the search of points of interaction between the enzyme and the relevant biochemical process. This interaction may be observed (1) by changes in the biological system as the activity of the enzyme is affected by inhibitors or activators, or (2) by finding factors that simultaneously affect both the function of the biological system and the activity of the enzyme. The relationship between the photochemical activity of PSII (measured at pH 5.5 and 6.5) and CA activity (measured at pH 8.3) had been shown earlier (Shitov et al., 2011, 2018), using first method, only for CO_2_ hydration, but not for HCO_3_^-^ dehydration. Here, at pH 6.5, we demonstrate, using the second method, a pronounced relationship between the photochemical activity of PSII and its HCO_3_^-^ dehydration activity by the following observations: (**i**) HCO_3_^-^ dehydration activity reaches its maximum at pH 6.5 when we go up from pH 5.5 (Lu and Stemler, 2002) or go down from pH 6.7 (Moskvin et al., 2004) (increasing by ∼650 and by ∼3,550 times, respectively), in parallel to results on O_2_ evolution (Schiller and Dau, 2000); (**ii**) at pH 6.5, Chl/RCII ratio in BBY preps (Table 1, line 8) affects, in the same way, the maximal quantum yield of photochemical reactions (Table 1, line 5), photosynthetic O_2_-evolution expressed in μmol O_2_·(mole RCII)^-1^·h^-1^ (Table 1, line 7) as it does the dehydration CA activity (Table 1, line 11). Thus, pH 6.5 implies as the optimum pH for both photochemical and CA activities in PSII. Further, we will discuss: (**1**) the main factor that could really affect both these activities in PSII (i.e. if pH 6.5 is really this factor); (**2**) the site, where the relationship between these activities occurs; and (**3**) the function of the CA activity in PSII.

### 3.1. The main factor that simultaneously influences the photochemical and CA activities of PSII

In this work, we have experimentally shown that CA activity increases by ∼8 times when the pH is decreased from 8.3 (Table 1, line 1) to 6.5 (Table 1, line 10). However, the conditions of the hydration reaction (pH 8.3) differed from the conditions of the dehydration reaction (pH 6.5) not only in pH, but also in other factors: the concentration of the substrate, the concentration of the buffer, the ionic strength and the temperature, used in the experiments (see “Materials and methods”). It should be noted that either of these factors, and not just pH, could cause the increase of CA activity. Here we have analyzed the influence of these factors on the increase of CA activity in our experiments to examine if the pH 6.5 is the main factor that simultaneously influences the photochemical and CA activities of PSII.

#### The effect of buffer composition

In our experiments, measuring dehydration reaction, we decreased the concentration of the buffer and its ionic strength by 3 and 50 times, respectively, as compared to the hydration reaction. Using CAII, it has been established that the decrease of both the buffer concentration (Nielsen and Fago, 2015) (and, consequently, of the substrate concentration) and the ionic strength (Kernohan, 1964) in the medium, results in the decrease of the rate of the enzymatic reaction. Taking into account that CA activity of PSII have enzymatic nature with the same mechanism of reactions (see “Results” section 2.3.) as that of CAII used in works (Nielsen and Fago, 2015) and (Kernohan, 1964), we should expect a decrease of the dehydration activity of PSII in comparison with the hydration one. But, on the contrary, we observed a large increase of the dehydration activity by ∼ 8 times (Table 1, lines 1 and 10). This result indicates that the composition of the buffer is not the reason of the increase of CA activity of PSII when we measured it at pH 6.5.

#### The effect of the temperature

In our measurements on PSII, we increased the temperature from 1-2 °C (the hydration reaction) to 25 °C (the dehydration reaction), i.e. by 23-24 °C (see “Materials and Methods”). Since the enzymatic nature of CA activity in PSII has been proved in our work (see “Results” section 2.3.), and the mechanism of catalysis appears to be the same as in other CAs, we expect a similar increase of CA activity under a similar increase of the temperature for both PSII and CAII. It is known that when the temperature is increased from 0 ^о^C to 37 °C (i.e. by 37 °C), one observes an increase of k_cat_ (and, consequently, of the CA activity) by no more than 2.9–5 times for the dehydration activity of α-CAs (Sanyal and Maren, 1981). Consequently, the rise of the temperature from 1-2 °C to 25 °C (i.e. by ∼24 °C) is expected to result in the increase of CA activity of PSII by no more than 1.5-2.5 times. In fact, this increase is much more pronounced (∼8 times (Table 1, lines 1 and 10)) in our data, and, therefore, the temperature is not the main factor in increasing CA activity. We suggest that pH 6.5 is the only major factor that could additionally (∼4 times) increase the CA activity in our experiments. Thus, we assume that the use of pH 6.5 (among all the mentioned factors) is the main reason of the observed increased CA activity of PSII.

We note that the influence of pH on both CA activity and O_2_ evolution is similar. I.e. the dehydration activity, has its maxima at pH 6.5 (Table 1, line 10), exceeding the activity at pH 5.5, by ∼650 times (see (Lu and Stemler, 2002)), and at pH 7.5, by ∼13 times (cf. k_cat_ in Table 1 (line 17) with k_2_ in Ref. (McConnell et al., 2007)). Further, the rate of O_2_ evolution is known to be maximal at pH 6.0 - 6.5, exceeding the activity at pH 5.5, by ∼1.5 times, and at pH 7.5, by ∼2.2 times (Schiller and Dau, 2000). The same manner of pH effect indicate that this factor can influence on both activities simultaneously. Consequently, pH is the key relating O_2_ evolution to CA activity. In addition, the above observation can indicate on the same site of O_2_ evolution and HCO_3_^-^ dehydration in PSII. Since it is difficult to separate which side of PSII (acceptor or donor) is affected by the change of pH (Schiller and Dau, 2000), CA activity may be participating in reactions on both sides of PSII. Further we will discuss were HCO_3_^-^ dehydration occurs in fact.

### 3.2. The site of the relationship between O_2_ evolution and HCO_3_^-^ dehydration

The number and the nature of active centers of CA in PSII is a complicated issue, since the available data about CA activity of different components of PSII is controversial. Several authors (Lu et al., 2005; Ignatova et al., 2006; Lu and Stemler, 2007; Rudenko et al., 2007; Ignatova et al., 2011) have proposed the presence of two locations of CA activity in PSII. This proposal is in agreement with our above assumption that CA activity may be associated with both sides of PSII. However, there is no agreement on the nature of CA activity centers and their locations in PSII, but many authors associate the CA activity of PSII with the Core-complex in this photosystem (Rudenko et al., 2015). To understand the nature of active centers we should: (1) determine how deep CA reactions occur in PSII and, after that, (2) find suitable places for these reactions in PSII.

#### 3.2.1. The characteristics of CA activity, showing that CA reactions occur deep inside the hydrophobic part of PSII

It is well known, that a metal ion (predominantly the Zn) catalyzes CA reactions in all CAs, locating at the bottom of the cavity of the active center of these enzymes (Supuran, 2008). Consequently, we can expect that any protein will have an active center with a metal ion in its bottom, if it enzymatically accelerates CA reactions (having the same mechanism of them). The CA of PSII, in fact, accelerates enzymatically HCO_3_^-^ dehydration and has the same mechanism of catalysis as bovine α-CAII (for details see section 2.3. of “Results”), and, therefore, the active center of CA in PSII should also contain metal ions in its bottom.

We have interested, may the depth of active centers of CAs (i.e. the depth of location of metal ion) correlate with catalytic properties, in principle? Our interest was caused by the publication of Aggarwal et al. (Aggarwal et al., 2013), in which the authors have proposed that the differences in catalytic efficiency among human α-CAs may be attributed to the varying speed of proton shuttling from the active center to the bulk solution via the network of amino acid residues. The proton shuttling speed have found to be the rate limiting step (which is reflected in the steady state parameter k_cat_) in the mechanism of catalysis for α-, β- and γ-CAs (Zimmerman et al., 2007, 2010; Supuran, 2008). So, the k_cat_ (being per se the frequency of the reaction) should inversely depend on the time of proton transfer (the rate limiting step of this reaction), because the frequency inversely depends on the time. Consequently, the time of proton transfer should directly depend on the length of the network of amino acid residues (through which the proton passes) in CAs, because the time is directly proportional to the distance. The length of the network of amino acid residues in turn, could directly depend on the depth of the cavity of an active center, i.e. on the distance between the entrance of the cavity and a metal ion in its bottom. Thus, we assume that k_cat_ could inversely depend on the depth of the cavity of an active center, which indicate that catalytic properties, in principle, may correlate with the depth of active centers, at least, in α-CAs.

To understand: whether the above assumption is true for PSII? we must first determine whether this assumption is true for all known CAs, which have the same mechanism of catalysis? For this we have determined the depth of active centers (using X-ray structures) for three CAs (Fig. 10, panel A), having the same mechanism of catalysis (Zimmerman et al., 2007): for human α-CAII (Avvaru et al., 2010), which is very close to bovine α-CAII (studied in this work) in structure and catalytic properties; for β-CA from *Pisum sativum* (Kimber and Pai, 2000); and for γ-CA CAM from *Methanosarcina thermophila* (Iverson et al., 2000). We chose these CAs since we know well their catalytic properties: human α-CAII has k_cat_ = 0.5·10^5^ - 2.26·10^5^ sec^-1^ at pH 6.5 - 7.0 and K_m_= 8.2 - 9 mM (Khalifah, 1971; McIntosh, 1968); β-CA from *Pisum sativum* has k_cat_ = 2.30·10^5^ sec^-1^ at pH 8.5-9.0 (Smith and Ferry, 2000) and K_m_= 52 mM (Johansson and Forsman, 1993); and γ-CA CAM has k_cat_ = 2.31·10^5^ sec^-1^ at pH 7.5 (Zimmerman et al., 2010; MacAuley et al., 2009) and K_m_= 59 mM (MacAuley et al., 2009). Unfortunately, the k_cat_ of β-CA from *Pisum sativum* is not determined at pH 6.5 – 7.0 yet, but it is well-known that with decreasing pH, k_cat_ of CAs also decreases (Gibbons and Edsall, 1964; Khalifah, 1971), therefore, we can expect lower k_cat_ of the β-CA at pH 6.5 – 7.0. We found that k_cat_ (at pH close to 7.0) actually increases in the row α-CAII → β-CA → γ-CA, when the depth of their reaction center decreases in this row (Fig. 10, panel A) (the last finding is confirmed by data in Ref. (Capasso and Supuran, 2015)). Thus, we concluded that, among α-, β- and γ-CAs (having the same mechanism of catalysis), the value of k_cat_ is inversely proportional to the depth of the active center of CAs.

**Figure 10.**
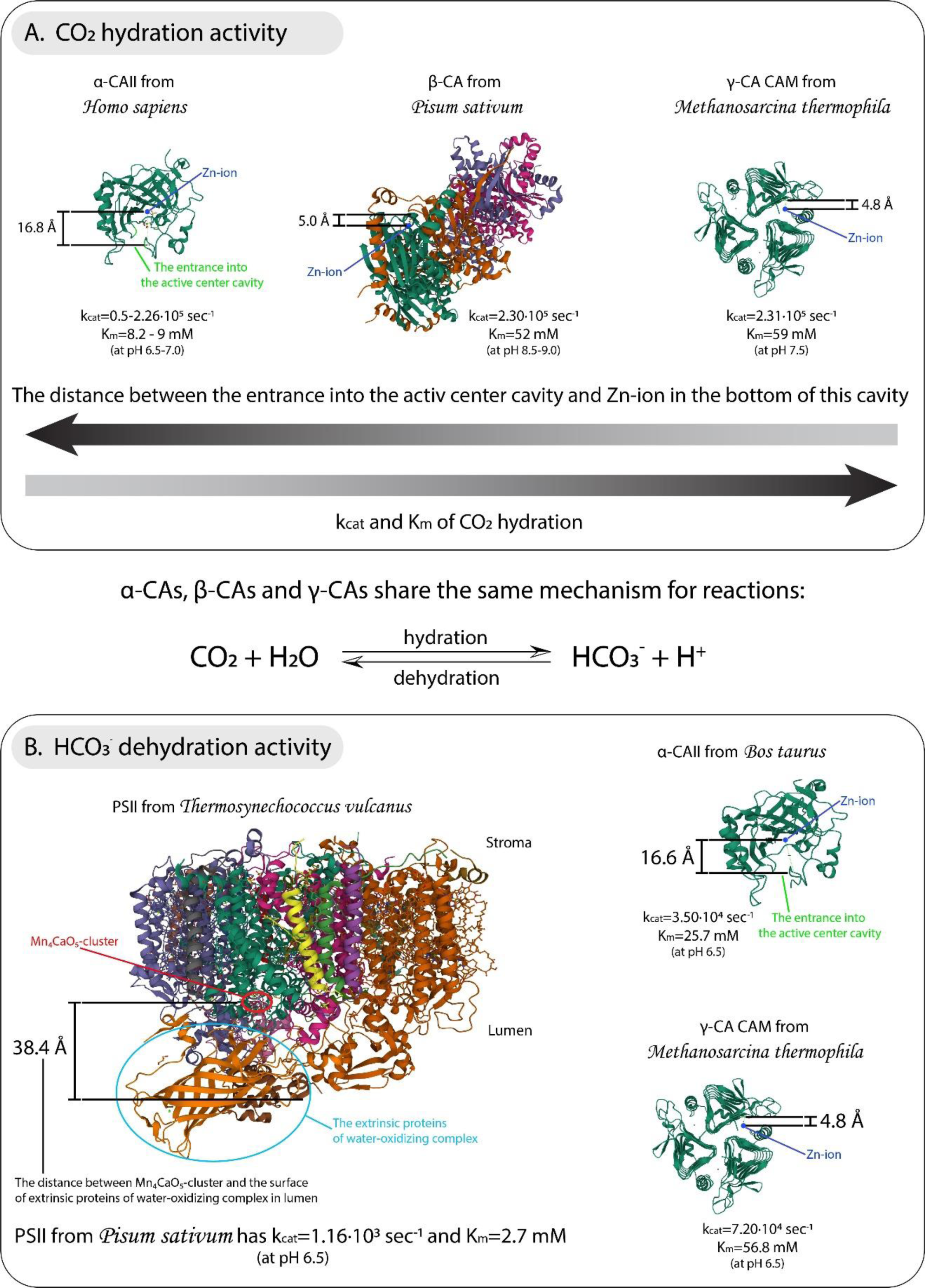
The interrelation of the values of k_cat_ and K_m_ with the distance between the entrance into the active center cavity and a metal ion in the bottom of this cavity in CAs. Panel A shows how k_cat_ and K_m_ for CO_2_ hydration depends on this distance in α-, β- and γ-CAs. Panel B shows that k_cat_ and K_m_ for HCO_3_^-^ dehydration depend, by the same manner as for CO_2_ hydration, on this distance in α- and γ-CAs. Here we use K_m_ and k_cat_ obtained in current work for PSII from *Pisum sativum* (Table 1, lines 12 and 17, respectively) and for α-CAII from bovine erythrocytes (Table 2, column 2). We have created all these images by Mol* Viewer, doi: 10.1093/nar/gkab314, RCSB PDB. To analyze structures we used the following PDB identifiers (the associated publications we have cited in the text): α-CAII from human erythrocytes – 3KS3; β-CA from *Pisum sativum* (from the stroma of chloroplasts) - 1EKJ; γ-CA CAM from *Methanosarcina thermophila* - 1QRG; PSII from *Thermosynechococcus vulcanus* - 4UB6; α-CAII from bovine erythrocytes - 1V9E.

In addition, we noted that K_m_ depend on the depth of the active center by the same way as k_cat_ (Fig. 10, panel A). Since K_m_ is independent of pH (Khalifah, 1971) and the dependence of this parameter on the depth of active center is more illustrative, than that of k_cat_, we will use K_m_, along with k_cat_, to understand how deep CA reactions occur in PSII.

We have obtained kinetic parameters of CA activity in PSII only for HCO_3_^-^ dehydration, but above (Fig. 10, panel A) we used these parameters for CO_2_ hydration, because k_cat_ for HCO_3_^-^ dehydration is unknown for β-CAs yet. Thereby the question arises: is the rule, established for the hydration reaction (Fig. 10, panel A), valid for the dehydration reaction? It is well known that the mechanism of catalysis of CO_2_ hydration and HCO_3_^-^ dehydration is the same in all CAs (at least for α-, β- and γ-CAs) (Zimmerman et al., 2007). Consequently, we can expect the same correlation of k_cat_ and K_m_ with the depth of active center also for HCO_3_^-^ dehydration for all CAs among these three classes. It is true, because bovine CAII, possessing k_cat_ = 3.5·10^4^ sec^-1^ and K_m_ = 25.7 mM (Table 2, column 2) for HCO_3_^-^ dehydration at pH 6.5, has more deep active center (∼ 16 Å) than that of γ-CA CAM (∼ 4.8 Å), possessing higher k_cat_ = 7.2·10^4^ sec^-1^ and K_m_ = 56.8 mM (Alber et al., 1999) for HCO_3_^-^ dehydration at pH 6.5 (Fig. 10, panel B). Thus, we conclude that, k_cat_ and K_m_ are inversely proportional to the depth of the active center for HCO_3_^-^ dehydration as well, and this rule is universal for α-, β- and γ-CAs.

Based on above conclusion, we could expect that the catalytic center (the metal ion) of CA locates more deep inside the PSII complex than it locates in other CAs, because the CA activity of PSII, having the same mechanism of catalysis as other CAs, possesses lower k_cat_ as well as lower K_m_ for HCO_3_^-^ dehydration (Table 1, line 12), than that of CAII (Table 2, column 2) and of γ-CA CAM (Alber et al., 1999). To test this hypothesis, we should: (1) chose the metal ion in the structure of PSII that may serve as the possible catalytic center of CA activity; and (2) determine the distance between this metal ion and the closest external surface of PSII complex. Taking into account that CA activity is significant for the function of the donor side of PSII (Shitov et al., 2011, 2018) and that this side contains Mn ions, we chose these ions as possible catalytic center of CA activity in PSII. Consequently, we chose the lumenal surface of the proteins of WOC as the external surface of PSII, because this surface is closest to the Mn_4_CaO_5_- cluster. To determine the distance between the lumenal surface of PSII and Mn ions, we used the X-ray structure of PSII. Since sufficiently detailed crystal structure is not obtained for PSII from higher plants yet, we used the structure of PSII from *Thermosynechococcus vulcanus* (Suga et al., 2015). The distance between the lumenal surface of PSII and Mn_4_CaO_5_-cluster was found to be 30.2 – 38.4 Å, which, really, is ∼2 times more than that (Fig. 10, panel B) in bovine α-CAII (X-ray structure of this CA have described in Ref. (Saito et al., 2004)). Moreover, this distance is more than 6 times larger as compared to γ-CA CAM (Fig. 10, panel B) and β-CA from *Pisum sativum* (Fig. 10, panel A). Thus, we proved that the active center of CA in PSII locates more deep than that in other α-, β- and γ-CAs.

We speculate that the distance between the lumenal surface of the proteins of WOC and Mn_4_CaO_5_-cluster is even more in PSII from higher plants (more than ∼38.4 Å) than in PSII from cyanobacteria, since the proteins of WOC (with the exception of PsbO) in PSII from higher plants have a mass (and therefore a size) more than that of PSII from cyanobacteria. Thus, we proved that the active center of CA activity in PSII from *Pisum sativum* locates deep inside the hydrophobic part of PSII complex (more deep than in other CAs), which causes low K_m_ and relatively low k_cat_ for HCO_3_^-^ dehydration.

#### 3.2.2. Suitable places for CA activity in PSII

Summarizing all the said above, the location of active center of the CA in PSII should be characterized by three criteria: 1) it should locate deep inside the hydrophobic part of PSII complex; 2) it should be near the metal ion in the structure of PSII; 3) it should be a part of Core-complex. Khristin et al. (Khristin et al., 2004) were the first who suggested the location of active center of CA inside the hydrophobic part of the PSII complex. Moreover, they also proposed that: «the Core-complex component possessing CA activity must have an active metal containing center, the structure of which resembles the structure of the known CAs. However, the metal is not obligatory to be zinc»… «Thus, it is not improbable that manganese within the water-oxidizing complex may be involved in the CA activity». Thus, one of the Mn ions of the Mn_4_CaO_5_-cluster, on the electron donor side, may be the one out of two active centers of CA activity in PSII. Below we present different arguments for Mn as the active center of CA in PSII.

##### 3.2.2.1. Mn as one of the active centers of CA activity in PSII

These two facts indirectly testify in favor of Mn: **(i)** Mn can replace Zn in the active center of some CAs, while maintaining their catalytic function (Tripp et al., 2004); **(ii)** it locates deep inside the hydrophobic part of PSII complex (Fig. 10, panel B). Below we will present the properties of CA activity, obtained in this work, that directly confirm the participation of Mn in the acceleration of CA reactions in PSII.

**(iii)** Taking into account that pH simultaneously, and in the same manner, affects both O_2_ evolution and HCO_3_^-^ dehydration (see section 3.1. of “Discussion”) and that O_2_ evolution occurs on the electron donor side of PSII, we propose that there is some event (on the donor side), caused by the change of pH, that simultaneously changes these activities. The release of Mn from the Mn_4_CaO_5_-cluster as well as the release of extrinsic proteins from the donor side of PSII, caused by the increase of pH (Schiller and Dau, 2000; Kuwabara and Murata, 1982; Klimov et al., 1982), may be this event, that results in the decrease of both O_2_ evolution (Schiller and Dau, 2000) and the hydration activity (Table 1, line 1) at pH 8.3, as well as in the lowering of the dehydration activity, described in Ref. (McConnell et al., 2007). We assume that even the destabilization of WOC, specifically, of the Mn_4_CaO_5_-cluster, affects the CA activity of PSII. The (electron) donor side of PSII, in general, and Mn_4_CaO_5_-cluster, in particular, are much more stable at pH 6.5 (showing maximal O_2_ evolution (Schiller and Dau, 2000)) and, therefore, this pH promotes, we suggest, higher dehydration activity (Table 1, line 10). Thus, we conclude that HCO_3_^-^ dehydration may be connected with the state of Mn_4_CaO_5_-cluster, showing the key role of Mn in CA activity of PSII. Below we present facts that additionally confirm this conclusion.
**(iv)** The inhibition of both CA activity and electron transfer by sulfonamides (in particular, by EA and AA) on the donor side of PSII (Shitov et al., 2011, 2018) also indicates the involvement of Mn in CA activity function, because it is well known that the metal ion in active center of all CAs directly binds sulfonamides, which decreases the rate of CA reactions (Supuran, 2008). Below we will discuss the latter fact in more detail.

If we propose one of the Mn of Mn_4_CaO_5_-cluster as the active center of CA activity in PSII, then sulfonamides EA and AA (which often used in investigations of this activity (Ignatova et al., 2006; Rudenko et al., 2007; Shitov et al., 2009, 2011, 2018)) can bind to Mn, inhibiting CA activity of PSII. Since Mn_4_CaO_5_-cluster locates deep inside the hydrophobic part of PSII, we propose that the efficiency of the inhibition of CA activity by sulfonamides may depend on the lipophilic (hydrophobic) properties of the specific inhibitor, because more lipophilic inhibitor penetrates more easily through the hydrophobic regions of protein(s) to the metal containing active center and, in turn, inhibits CA activity more efficiently. We (Table 1, and (Shitov et al., 2009, 2011, 2018)) and others (Ignatova et al., 2006; Rudenko et al., 2007; Moskvin et al., 2004) have shown that the efficiency of the inhibition of CA activity in PSII by sulfonamides really depend on the lipophilic properties of the inhibitor. We demonstrated (Table 1, lines 3 and 4) that EA, having lipophilic properties (Moskvin et al., 2004), is ∼2 times more effective than AA, having hydrophilic properties (Moskvin et al., 2004), which proves at least that the active center of CA locates inside the hydrophobic part of PSII.

We propose that the proteins of WOC (PsbO, PsbP and PsbQ) may serve as a barrier, which prevents the penetration of hydrophilic AA into the PSII complex. After removing this barrier, we can expect the increase of the efficiency of inhibition of CA activity by AA. We confirm this suggestion by our early results (Shitov et al., 2009), where we have shown that after complete removal of PsbO, PsbP and PsbQ from BBY, the inhibition of CA activity by AA increased, becoming equal to that of EA (Fig. 11). Thus, the active center of CA in PSII should be located behind the proteins of WOC (see the diagram in the Fig. 11). Therefore, we concluded that one of the Mn ions of Mn_4_CaO_5_-cluster actually can belong to the active center of CA in PSII, since Mn locates behind the proteins of WOC and inside the hydrophobic part of PSII.

**Figure 11.**
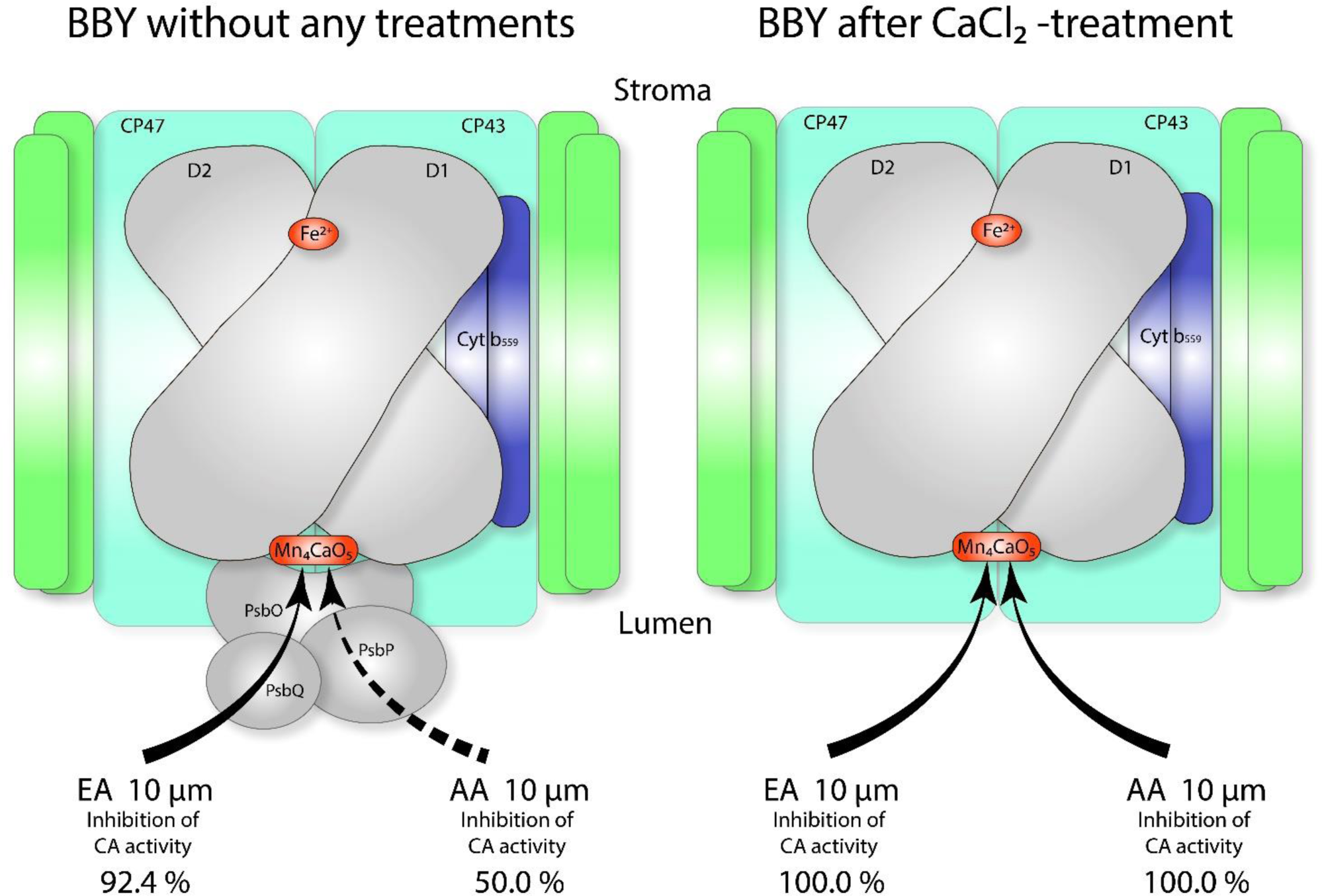
The extrinsic proteins of water-oxidizing complex as a barrier for the penetration of acetazolamide inside the hydrophobic part of PSII complex. On the left we show how lipophilic EA and hydrophilic AA inhibit the hydration CA activity of BBY, which contains the extrinsic proteins of WOC PsbO, PsbP and PsbQ. We have taken the data of inhibition in the Table 1. On the right we present how EA and AA inhibit the hydration CA activity of BBY, depleted of the extrinsic proteins. The proteins PsbO, PsbP and PsbQ were completely removed from BBY by CaCl_2_-treatment, saving Mn_4_CaO_5_-cluster in the composition of obtained preparation (we have taken these results from our earlier work, cited in the section 3.2.2.1. of “Discussion”). D1 and D2 are the reaction center proteins, Cyt b_559_ is a cytochrome b_559_. CP43 and CP47 are Chl-protein complexes of 43 and 47 kDa that form the inner antenna system of PSII; PsbO (33 kDa), PsbP (23 kDa) and PsbQ (16 kDa) are extrinsic proteins that stabilize and optimize the water-oxidizing complex and its function. Fe^2+^ is the non-heme Fe, Mn_4_CaO_5_ is the manganese-oxygen-calcium cluster.

##### 3.2.2.2. Non-heme Fe as second active center of CA activity in PSII

Working properly Core of PSII contains not only Mn, but also non-heme Fe on the electron acceptor side of this complex. Thus, non-heme Fe, on the electron acceptor side, may be the second out of two active centers of CA activity in PSII. Here we present some arguments, confirming our suggestion: **(i)** Fe can act as a catalytic center of CA activity in γ-CA CAM (in anaerobic conditions) and, moreover, this CA possesses higher catalytic efficiency than the same CA, containing Zn, instead of Fe (Tripp et al., 2004; Alber et al., 1999; MacAuley et al., 2009; Zimmerman et al., 2010). **(ii)** Fe, similar to Mn, also locates deep inside the hydrophobic part of PSII complex. We present the model of location of active centers of CA activity in PSII on the scheme Fig. 12 (A). **(iii)** BBY preparations (depleted of PsbO, PsbP and PsbQ (see ref. (Shitov et al., 2009)), even when they are depleted of Mn_4_CaO_5_-cluster (but having non-heme Fe), still show CA activity. **(iv)** Non-heme Fe have found to bind HCO_3_^-^ at the electron acceptor side of PSII (Brinkert et al., 2016; Ferreira, 2004; Suga et al., 2015). In spite of this arguments, the function of non-heme Fe as the active center of CA in PSII needs additional investigations. Nevertheless, the existence of 2 active centers of CA (the nonheme Fe on the electron acceptor side of PSII and the Mn atoms in Mn_4_CaO_5_-cluster on the electron donor side of PSII), in general, is consistent with all the experimentally obtained results (Khristin et al., 2004; Lu et al., 2005; Lu and Stemler, 2002, 2007; McConnell et al., 2007; Ignatova et al., 2006, 2011; Shitov et al., 2011, 2018). Since, HCO_3_^-^ dehydration have found to be associated with the reactions, occurring on the electron donor side of PSII (see previous section), further we will discuss the function of CA activity on this side of PSII.

**Figure 12.**
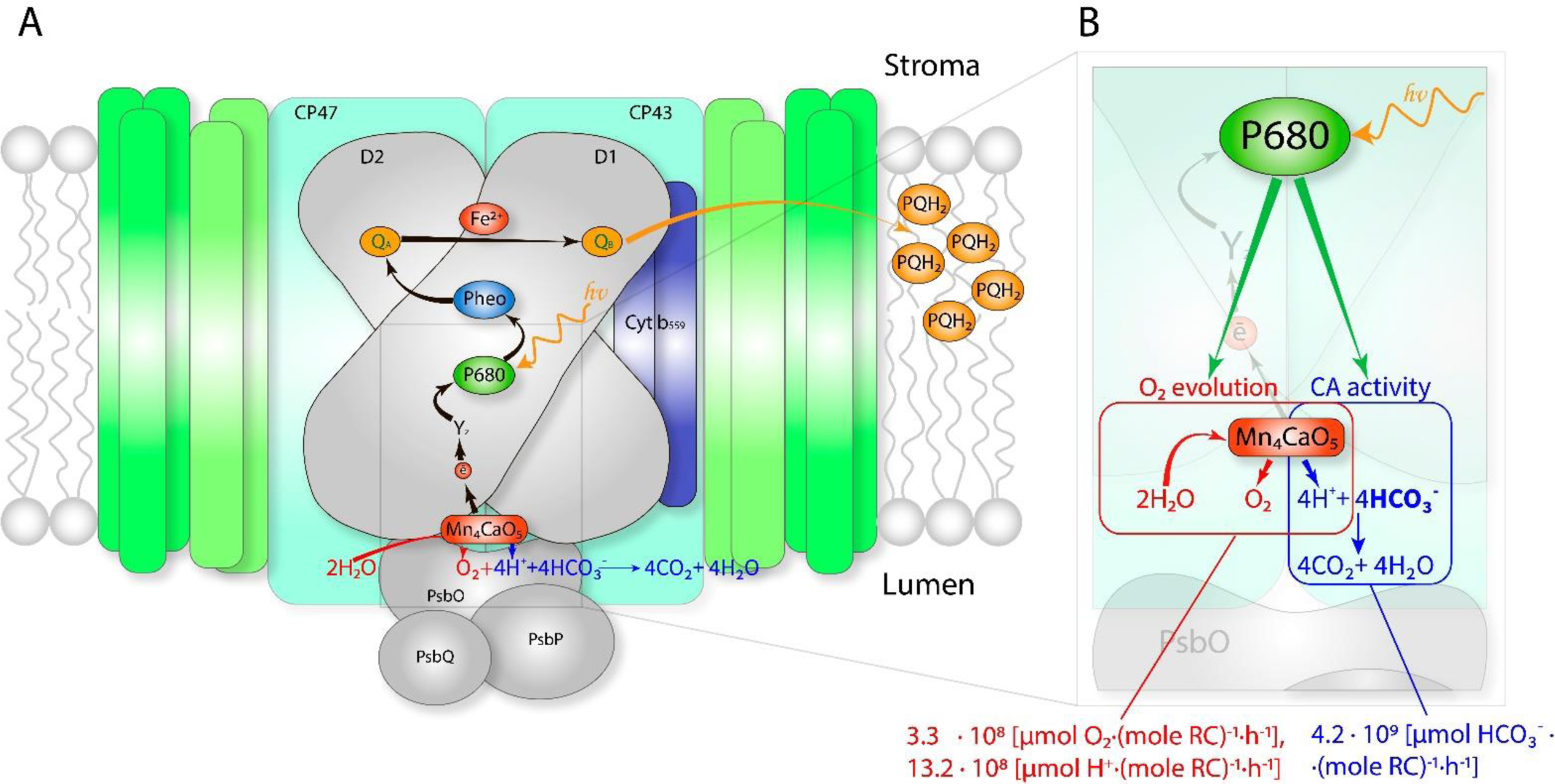
Probable sources of CA activity and its of function on the electron donor side of PSII. The sources of CA activity in PSII from higher plants are shown in red figures in panel (A), the proposed function of this activity on the electron donor side is schematically presented in panel (B). D1 and D2 are the reaction center proteins, Cyt b_559_ is a dimeric protein that contains the redox active cytochrome b_559_ (these 4 subunits forms RCII of PSII). CP43 and CP47 are Chl-protein complexes of 43 and 47 kDa that form the inner antenna system of PSII; PsbO (33 kDa), PsbP (23 kDa) and PsbQ (16 kDa) are extrinsic proteins that stabilize and optimize the water-oxidizing complex and its function. Fe^2+^ is the non-heme Fe, Mn_4_CaO_5_ is the manganese-oxygen-calcium cluster. P680 is a pair of Chl; Pheo, pheophytin on D1, is the primary electron acceptor; Q_A_ (on D2), bound plastoquinone; Q_B_ (on D1), exchangeable plastoquinone; Yz (on D1) is the redox active tyrosine residue in PSII and PQH_2_, mobile plastoquinone molecules in the membrane. Black arrows show electron transfer pathway through PSII complex. Blue arrows indicate the movement and transformation of substrates of CA reactions. Red arrows determine the movement and transformation of the substrate (H_2_O) and the release of the product (O_2_) of water oxidation reaction. Green arrows present that the charge separation act in P680 triggers both O_2_ evolution and CA reaction. Orange arrow show the way of the plastoquinol moving into plastoquinone pool inside the thylakoid membrane. The intersection of the red and blue rectangles (in the panel (B)) indicates the interaction point between the O_2_ evolution and CA reaction. This point seems to be the protons that released during water oxidation and HCO_3_^-^ ions (which accept the released protons).

### 3.3. The function of CA activity on the electron donor side of PSII

The function of CA activity in PSII may be equally related to the catalysis of both the hydration and dehydration reactions (see Eqs 2 and 3). The same dependence of both photochemical and dehydration activities on Chl/RCII ratio (as well as almost the same dependence of these activities on pH (Table 1, lines 1 and 10 and Ref. (Schiller and Dau, 2000)) implies that one common process may be involved in performing both these activities. This process may be a chemical and/or a photochemical reaction, or both. The only one photochemical process, the charge separation in the P680 of RCII, appears to be the key reaction, since the amount of charge separation directly depends on the number of molecules of Chl bound per RCII (and, subsequently, its efficiency directly depends on the number of quanta captured by these Chls). This suggestion is supported by our results, where one active center of CA was found to be related to one photochemical RCII in PSII (Fig. 7). Thus, we speculate that the charge separation act, triggering water oxidation, may also act as a trigger of CA reaction in PSII (see the model in Fig. 12 (B)).

Analyzing above assumption, the question arises: Is the charge separation the only one to trigger the process for both CA activity and O_2_ evolution, or is there an intermediate process that connects the charge separation and both the activities? We note that K_m_ = 2.7 mM, obtained here, for HCO_3_^-^ dehydration, is close to this parameter (1 and 1.4 mM) obtained for the bicarbonate effect on the O_2_ evolution by Khanna et al. (Khanna et al., 1977) and by vanRensen and Vermaas (van Rensen and Vermaas, 1981; van Rensen and Klimov, 2005). Close values of K_m_ for the photochemical activity and for the HCO_3_^-^ dehydration suggest a common mechanism of action of HCO_3_^-^ in both the processes. Accepting the hypothesis that HCO_3_^-^ is a mobile carrier involved in the removal of H^+^s from Mn_4_CaO_5_-cluster (Shutova et al., 2008; Koroidov et al., 2014; Shevela et al., 2020), we suggest that the protons, released during water oxidation, are the link between O_2_ evolution and CA activity via the reaction (Eq. 3) (see Fig. 12 (B)). Consequently, the reaction (Eq. 3) is suggested to be the common mechanism, combining these two reactions.

Above conclusion is supported by the following experiments. On the one hand, HCO_3_^-^, promoting the consumption of H^+^ (via the reaction (Eq. 3)), supports O_2_ evolution in PSII, as shown for the green alga *Chlamydomonas reinhardtii* (Shutova et al., 2008) and for spinach, *Spinacea oleracea* (Koroidov et al., 2014). On the other hand, CA activity (i.e. the acceleration of the reaction (Eq. 3) described here, in particular) was found to be significant for maximal photochemical activity (O_2_ evolution, in particular) in PSII from *C. reinhardtii* (Terentyev et al., 2019; Shukshina and Terentyev, 2021) and from pea, *Pisum sativum* (Shitov et al., 2011, 2018). Thus, we propose that the charge separation in P680 of RCII, ultimately, triggers the reaction (Eq. 3), which, being catalyzed by CA activity, further directly participates in O_2_ evolution on the electron donor side of PSII (see scheme in Fig. 12 (B).

This hypothesis may be correct, if the products of reaction (Eq. 3) can be shown to accumulate at a rate that exceeds O_2_ evolution at least by 4 times, since the evolution of 1 mole of O_2_ is accompanied by the formation of 4 moles of H^+^s (Fig. 12). It is important to note that although Koroidov et al. (Koroidov et al., 2014) have experimentally shown the formation of CO_2_ (one of the products of the reaction (Eq. 3)) that is associated with the formation of O_2_ in the process of photosynthetic water oxidation in BBY preps from spinach, yet the rate of CO_2_ formation was much less than the rate of O_2_ evolution (Koroidov et al., 2014). However, in the work, presented here, we have found that the k_cat_ of HCO_3_^-^ dehydration in PSII (Table 1, line 17) is ∼13 times greater than the rate of photosynthetic O_2_ evolution (Table 1, line 7), and it exceeds by ∼3.2 times the formation of protons (Fig. 12). Therefore, we conclude that the rate of HCO_3_^-^ dehydration (accelerated enzymatically by CA activity) is sufficient for effective H^+^ consumption from the Mn_4_CaO_5_-cluster in the process of photosynthetic water oxidation.

So, we experimentally confirm here the function of CA activity as the catalyst of the reaction (Eq. 3), in the process of photosynthetic water oxidation on the electron donor side of PSII. This does not preclude CA activity (and its function) also on the electron acceptor side of PSII. Thus, the CA activity is an integral part of the mechanism of function of PSII and determines not only the efficiency of photochemical reactions in this complex, but also the overall productivity of photosynthesis of plant cells (taking into account that PSII is the only source of electrons for the electron transfer chain in thylakoids).

## 4. Materials and Methods

### 4.1. Sample preparation and activity measurements

Purified CA from bovine erythrocytes (containing CAII isoform) was obtained from Sigma (USA) Cat. № C3934. Phoosysteem II (BBY) preparations were prepared following the method described in Ref. (Berthold et al., 1981), with modifications, as in Ref. (Schiller and Dau, 2000). These preparations were obtained from 14-day-old pea (*Pisum sativum*) leaves (from plants grown in the greenhouse).

The photosynthetic activity of the BBY preparations was measured by two methods. (1) The oxygen-evolving activity, as described in Ref. (Shitov et al., 2018). (2) The variable Chl fluorescence of PSII (or so-called “delta F”), as described in reference (Karacan et al., 2016). The total Chl content, in the preparations, was determined by the method of Porra et al. (Porra et al., 1989).

To prepare an extract of PSII pigments, BBY preparations (with 250 µg of Chl) were sedimented at 10,000 rpm for 3 min. The obtained pellet was diluted with ethanol (96%) to 600 µL. This mixture was shaken and then centrifuged at 12,000 rpm for 10 min, using a Mini Spin centrifuge (Eppendorf, Hamburg, Germany). A supernatant, containing the extract of pigments, was used for measurements (final Chl concentration, 4.2 µg·mL^-1^). Since the obtained precipitate had a very light color (without any green or yellow tint), this indicated that we had obtained full extraction of pigments from the PSII preparations. The Chl/RCII ratio was determined according to published methods (Kaminskaya et al., 2005; Terentyev et al., 2020).

### 4.2. The hydration and dehydration carbonic anhydrase activity measurements

The hydration activity for CA was measured, as described in reference (Wilbur and Anderson, 1948) with modifications, described by others (Shitov et al., 2009; Karacan et al., 2016). The details of the dehydration carbonic anhydrase activity were as follows. Pyranine fluorescence measurements were made using a Varian Cary Eclipse (Agilent Technologies, USA) fluorescence spectrophotometer; it was equipped with an RX.2000 Stopped-Flow Mixing Accessory (Applied Photophysics, United Kingdom). The stopped-flow apparatus was used in combination with a Pneumatic drive DA.1 to ensure uniformity of the mixing process. The mixing time was 8 ms. Chamber A contained a substrate – NaHCO_3_ ≥99.5% purity (Sigma, USA) in 2.0 mL of buffer A (0.5 mM Tricine-KOH) at pH 8.0. Chamber B contained CAII or BBY particles in 2.0 mL of buffer B (5 mM HEPES-KOH, 5.5 µM pyranine) at pH 6.5. We mixed the solutions kept in chambers A and B in a ratio of 1/1 at 3 Bars (43.5 psi) and at 25 °C. Pyranine fluorescence was measured at an emission wavelength of 512 nm (where changes are pH dependent); excitation light was 466 nm (Shingles and Moroney, 1997). All the slits of monochromator were set at 5 nm. The plots of the kinetics for pyranine fluorescence changes (which required smoothing of curves in some cases) and the fitting of Lineweaver-Burk plots, as well as statistical analyses, were carried out in Origin 6.5 software package (OriginLab, Northampton, Massachusetts, USA).

**Steady state initial rates** were calculated from the kinetic data (first 3-5 sec after HCO_3_^-^ addition), and expressed in three different units: relative units of fluorescence per second (rel.un.fl.·sec^-1^); pH·sec^-1^ and (mM H^+^)·sec^-1^; all the calculations were made as published in Refs (Khalifah, 1971; Steiner et al., 1975; Shingles and Moroney, 1997; Pocker and Bjorkquist, 1977a). The “Buffer factor” Q was calculated according to the following equation:

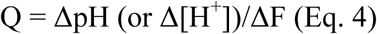

where, ΔpH is the change of pH per second, Δ[H^+^] is the change of concentration of protons per second and ΔF is pyranine fluorescence change per second. According to our calculations, Q was equal to 0.0109 ± 0.0001 pH·rel.un.fl.^-1^ and (2.34 ± 0.14)·10^-2^ mM ·rel.un.fl.^-1^, respectively. We calculated Δ[H^+^] through the HCO_3_^-^ concentrations (at the start and at the end at equilibrium of the reaction (Eq. 1)) which were calculated from the Henderson-Hasselbalch equation (pKa 6.48, 25 °C (Nielsen and Fago, 2015)) taking into account the concentration of added bicarbonate. These calculations were based on the equation Δ[H^+^] = Δ[HCO_3_^-^], presented in Ref. (Pocker and Bjorkquist, 1977a). For our measuring system, we found that Q remained constant over the entire range of HCO_3_^-^ concentration used. All pH measurements were made by a pH-meter (Orion 420Aplus; Thermo Electron Corporation, USA).

Taking into account the drop of pyranine fluorescence in the presence of BBY preps and based on a large number of measurements (more than 100) at 4.2 (µg Chl)·mL^-1^, we found the constant factor of this drop was equal to 1.71 ± 0.013. To compare properly the initial rate of the reaction in PSII with that of CAII, we have multiplied the results obtained on BBYs (the amount of pyranine fluorescence) by this factor. So, to keep the same level of the drop of pyranine fluorescence for all preparations, we used the Chl concentration as a basis for all measurements of dehydration CA activity in BBYs.

### 4.3. Calculation of carbonic anhydrase kinetic parameters (V_max_, K_m_, k_cat_ and k_cat_/K_m_)

Lineweaver-Burk plots (Lineweaver and Burk, 1934) (1/V versus 1/S) were used to determine V_max_ and K_m_, where, V is the initial velocity of catalyzed reaction, and S is the substrate (HCO_3_^-^) concentration. We used the obtained plots to derive their equations in the Origin program. Then we used the derived equations to calculate K_m_ and V_max_ values. k_cat_ was calculated according to the following equation

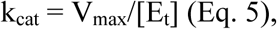

where, V_max_ is the maximal velocity expressed in both pH·sec^-1^ and (mM H^+^)·sec^-1^; [E_t_] – CAII concentration equaled 1.725·10^-5^ mM (0.5 µg·mL^-1^). For BBYs the [E_t_] is the RCII concentration, calculated separately for each PSII preparation (Table 1).

## Acknowledgments

We thank the late Professor V.V. Klimov (who initiated this work**^#^**) for discussions on the role of bicarbonate and CA activity in PSII function, M.S. Khristin for help in the search of suitable method of dehydration activity measurements, and A.A. Khorobrykh for help in the determination of RCII concentrations in the BBYs. Financial support was provided by Molecular and Cell Biology Programs of the Russian Academy of Sciences, by grants № 16-34-60028 and № 17-04-01011 from the Russian Foundation for Basic Research. The results presented in Table 1 were obtained with the support from the Ministry of Education and Science of the Russian Federation (theme AAAA-A17-117030110136-8). Govindjee thanks the staff of the Department of Plant Biology and of the Department of Biochemistry (University of Illinois at Urbana-Champaign) for constant support during the course of this research.

^#^ S. I. Allakhverdiev, S.K. Zharmukhamedov, M.V. Rodionova, V.A. Shuvalov, C. Dismukes, J-R.. Shen, J. Barber, G. Samuelsson and G. Govindjee : <u>Vyacheslav (Slava) Klimov (1945-2017): A scientist par</u> <u>excellence, a great human being, a friend, and a Renaissance man.</u> *Photosynth. Res*. **136**, 1-16 (2018).

## Author Contributions

A.S. designed and performed research, analyzed data, wrote the paper; V.T. performed research; G.G. wrote the paper.

## Contact information of authors

1. Alexandr V. Shitov; Phone: +7(4967)731966.
2. Vasily V. Terentyev; Phone: +7(4967)731966.
3. Govindjee Govindjee; Work Phone: 217-333-1794, Home Phone: 217-721-9954

